# Recruitment of MRE-11 to complex DNA damage is modulated by meiosis-specific chromosome organization

**DOI:** 10.1101/2020.06.26.174573

**Authors:** Kailey Harrell, Madison Day, Sarit Smolikove

## Abstract

DNA double-strand breaks (DSBs) are one of the most dangerous assaults on the genome, and yet their natural and programmed production are inherent to life. When DSBs arise close together (clustered) they are particularly deleterious, and their repair may require an altered form of the DNA damage response. Our understanding of how clustered DSBs are repaired in the germline is unknown. Using UV laser microirradiation, we examine early events in the repair of clustered DSBs in germ cells within whole, live, *Caenorhabditis elegans*. We use precise temporal resolution to show how the recruitment of MRE-11 to complex damage is regulated, and that clustered DNA damage can recruit proteins from various repair pathways. Abrogation of non-homologous end joining or COM-1 attenuates the recruitment of MRE-11 through distinct mechanisms. The synaptonemal complex plays both positive and negative regulatory roles in these mutant contexts. These findings together indicate that MRE-11 is regulated by modifying its accessibility to chromosomes.

## Introduction

Clustered DNA double-strand breaks (DSBs) are DSBs found in close proximity to each other (Sharma et al., 2016). These forms of DNA damage are particularly deleterious as they pose challenges to the DSB repair response. Studies in tissue culture have shown that repair of clustered DSBs involves a shift in DSB repair pathway utilized from non-homologous end joining (NHEJ) to homologous recombination (HR) (Wang et al., 2010). Whether a change in repair pathway choice occurs in cells that are already committed to HR is unknown. One of the tissues most overlooked in studies of repair of clustered DSBs is the germline, a complex tissue that generates the gametes of an organism through a specialized reductional division termed meiosis. The germline originates at a population of stem cells that proliferate mitotically and from which a fraction can differentiate and enter meiosis. Meiotic cells then undergo an extended meiotic prophase I in which pairs of homologous chromosomes associate with each other via crossovers and a proteinaceous structure called the synaptonemal complex (SC). These crossovers are necessary for the correct segregation of homologs at the first meiotic division to halve the complement of chromosomes and produce viable, haploid gametes. To ensure that these crossovers form, DSBs are intentionally induced during meiotic prophase I (Keeney, 2008). Despite their toxicity, DSBs play a crucial role in this meiotic context. Because DSBs are both necessary and hold incredible potential for mutagenesis, the accurate repair of DSBs is a tightly regulated process.

Programmed meiotic breaks are induced by the topoisomerase-like protein Spo11 (Keeney et al., 1997). Sae2/CtIP/Com-1 promotes the cleavage of the Spo11-DNA adduct by the Mre11-Rad50-Xrs2/Nbs1 (MRX/N) complex before the initiation of resection (Anand et al., 2016, Cannavo & Cejka, 2014). Breaks are resected to create single-stranded DNA (ssDNA) that is then bound by the RPA complex, followed by Rad51 (Wang & Haber, 2004). The SC forms between homologs to keep them close together upon the formation of DSBs and aids in the repair process (Hamer et al., 2008, Page & Hawley, 2004). This close association of homologs mediated by the SC, as well as the generation of ssDNA at the sites of Spo11-induced breaks, ensures that repair proceeds down the pathway of HR (Shibata et al., 2014). HR is generally considered to be an “error-free” method of repair, as it involves the use of a homologous repair template (Mehta & Haber, 2014). In the case of meiosis, the template employed for repair is the homologous chromosome, which aids in the establishment of a crossover between homologs and faithful segregation at the first meiotic division.

The covalent binding of Spo11 to generate intentional breaks precludes most other repair pathways from operating on these DSBs, since ends bounds by Spo11 cannot be ligated (Povirk, 2012). Removal of Spo11 creates ssDNA that is compatible with HR, but not with NHEJ. The NHEJ pathway is an error-prone repair pathway, associated with indels and rearrangements (Lieber et al., 2003). In mammalian meiotic cells and in yeast meiosis, NHEJ is also inhibited by the downregulation of its components inside of meiotic nuclei (Goedecke et al., 1999, Lee et al., 1999, Valencia et al., 2001). Nonetheless, it has been shown in some organisms that NHEJ can act at meiotic DSBs when HR is abrogated, indicating that NHEJ is not necessarily non-functional during meiosis (Yin & Smolikove, 2013, Yun & Kim, 2019). While the presence of a covalently-bound protein drives repair towards HR with programmed meiotic DSBs, it is unclear how regulation of repair pathways occurs at sites of exogenous damage in a meiotic context.

In addition to meiotic DSBs, the germline, like any other tissue, is exposed to other forms of DNA damage. This damage can be created by endogenous factors (such as collapsed replication forks) or exogenous factors (such as ionizing radiation). The repair of such damage has potential to be different than the repair of Spo11-induced breaks, either due to the type of break generated and/or the timing of break formation. One major difference between the types of breaks is the structure of the DSB ends. HR of Spo11-generated breaks requires Mre11/CtIP for end processing since this is the only nuclease that can remove the covalently bound Spo11 (Hartsuiker et al., 2009). However, breaks that do not contain adducts should not require the activity of Mre11/CtIP and other nucleases can promote resection independently of this complex (Mimitou & Symington, 2008). Unlike meiotic DSBs that form only at the entry to meiotic prophase, other forms of DSBs may occur outside this stage. This is particularly true for ionizing radiation which can occur at any stage of meiosis.

Laser microirradiation has been used primarily in tissue culture studies to induce DNA damage and study the *in vivo* recruitment of fluorescently tagged repair proteins (Lukas et al., 2004, Popuri et al., 2012, Rogakou et al., 1999). The type of damage induced by microirradiation is dependent on the wavelength used and the power utilized, and ranges from single-stranded nicks to double-stranded breaks (Kong et al., 2009, Lan et al., 2004). The range of damage induced, as well as the close proximity of many different types of DNA damage (creating clusters of damage), makes the type of damage induced by laser microirradiation complex. Studies performed in mammalian tissue culture cells indicate that clustered DSBs are more deleterious than spaced DSBs (Nickoloff et al., 2020). Moreover, in these cells repair pathway choice shifts from NHEJ to HR. We have shown that microirradiation attracts HR proteins, as do induced and spaced SPO-11 DSBs (Koury et al., 2018). However, it is still unknown if clustered DSBs in the germline attract NHEJ proteins. The transparent nematode *Caenorhabditis elegans* offers a unique opportunity to study exogenous DNA damage in the context of clustered DSBs within the germline of a whole, live organism.

A main component of DNA damage repair in the *C. elegans* germline is MRE-11. MRE-11 is the nucleolytic member of the MRN protein complex responsible for generation of ssDNA in *C. elegans* meiosis (Chin & Villeneuve, 2001). MRE-11, with the help of COM-1, removes covalently bound SPO-11 for processing of programmed meiotic breaks, and in doing so helps to ensure that HR factors can access the ssDNA and begin repair. These essential roles of SPO-11 removal and generation of ssDNA to begin HR make the MRN complex, specifically MRE-11, a key site of regulation for DSB repair pathway choice in the germline. As such we set out to characterize the recruitment kinetics of this protein *in vivo*. Here, we show that microirradiation induces similar levels of damage throughout the *C. elegans* germline that is distinct from apoptotic damage. Using this damage induction method, we show that MRE-11 is recruited to microirradiation-induced DNA damage as early as 10 seconds following microirradiation throughout the germline. MRE-11 can form both foci and clusters at this damage, likely indicating that the damage induced is complex. This is supported by the recruitment of factors from other repair pathways, namely the cKU-70/80 heterodimer which is involved in NHEJ. We show that cKU-70 does not affect the recruitment time of MRE-11 but influences the formation of clusters in mitotic and early meiotic zones of the germline. In our investigation into accessory factors of the MRN complex, we demonstrate that COM-1 enhances both the recruitment and the activity of MRE-11 at sites of complex DNA damage in the context of a fully formed SC.

## Results

### MRE-11 exhibits recruitment kinetics to microirradiation-induced breaks consistent with its role in meiosis

We previously showed that the UV laser microirradiation system can generate DSB clusters in individual germline nuclei of live *C. elegans* without increasing apoptosis or damaging adjacent germline nuclei that were not microirradiated (Koury et al., 2018). Our previous studies were done in the *spo-11* mutant background since both RPA-1 and RAD-51 localize to SPO-11 induced breaks, which complicates the analysis of their recruitment to DSBs in meiotic cells. Therefore, to test if HR proteins localize to microirradiation induced breaks and form clusters of DSBs in wild-type nuclei we followed MRE-11, a member of the MRN complex and the major nuclease that processes meiotic DSBs (Mimitou & Symington, 2009, Ogawa et al., 1995). Consistent with its role in formation and processing of meiotic DSBs, MRE-11 is found in all germline nuclei (as evidenced by its concentrated nuclear haze) but does not form foci (Fig 1A (left image) and Fig 1B) (Reichman et al., 2018). MRE-11’s mechanism of action suggests that it can be used as an early marker for DSB repair following DSB cluster formation in meiosis, as found in mammalian (mitotic) tissue culture studies (Shamanna et al., 2016, Suhasini et al., 2013). We have shown previously that MRE-11::GFP is expressed in all germline nuclei, but the variation in levels of nuclear localization throughout the germline was not examined (Fig 1B) (Reichman et al., 2018). This is important to examine since MRE-11 recruitment to microirradiation-induced breaks may be influenced by its relative nuclear concentration. To determine if the amount of MRE-11 protein localizing to nuclei is altered throughout meiotic progression we measured the fluorescent intensity of MRE-11::GFP throughout four regions of the germline primarily examined in this study. The intensity of MRE-11::GFP expression significantly increases with progression through the germline (2524 PMT, 3561 TZ, 6691 MP, and 7548 LP; Fig 1B). This is consistent with MRE-11’s transition from a redundant role (mitosis) to an obligatory role (meiosis) in the repair of DSBs in the germline.

**Figure 1.**
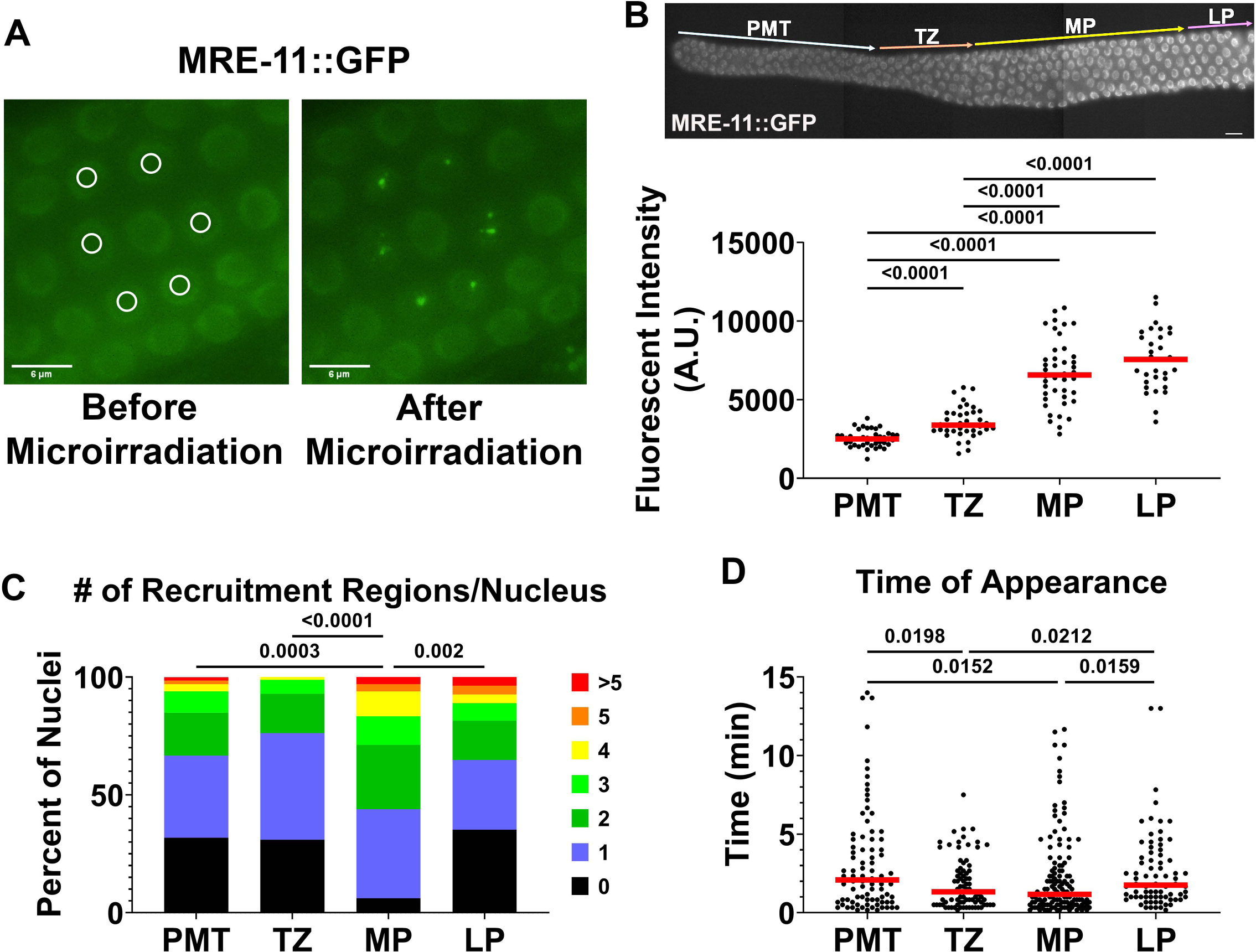
MRE-11 is recruited to microirradiation-induced DNA damage within the *C. elegans* germline. **A)** Example of microirradiation-induced foci in the *mre-11::gfp* strain. Left panel is MP nuclei before microirradiation, right panel is MP nuclei 4 minutes post-microirradiation. White circles in left panel indicate targeted nuclei. **B)** Top: A dissected gonad from an *mre-11::gfp* worm. Scale bar = 10µm. Bottom: Fluorescent intensity of MRE-11::GFP in the four main germline zones studied in this paper. Horizontal red line indicates median. Each point is an individual nucleus. Values were corrected to cytoplasmic background. **C)** Number of recruitment regions per nucleus after microirradiation. **D)** Time of recruitment region appearance in minutes after microirradiation. Each data point represents an individual region. Horizontal red line indicates median. For data sets with more than 2 groups (B, C, and D) Kruskal-Wallis was applied to determine significant differences between rank means, and if there was determination of significant differences, Mann-Whitney U-test was applied for each pairwise comparison.

To study MRE-11 recruitment to microirradiation-induced damage we had to determine the conditions of the live-imaging experiments (frames per minute) which affects the overall time of imaging. Acquiring live-imaging data is limited by photobleaching, thus longer time imaging allows later appearing foci to be detected, but needs to be performed at a lower resolution (less frames per minute). This prevents the detection of foci undergoing fast turnover and over-estimates time of appearance (*i*.*e*., foci appearing 10 seconds post-microirradiation will be recorded as appearing at 2 minutes, when acquisition is done every 2 minutes). Thus, the conditions of live-imaging experiments have tradeoffs, and results of the analyses performed is relative to the experimental conditions. In this manuscript we conducted live imaging experiments in a 15-minute window following microirradiation based on the rationale presented below. We previously demonstrated that upon microirradiation, RPA-1 and RAD-51 form foci at sites of damage ∼8 and ∼20 minutes post-microirradiation, respectively (Koury et al., 2018). First, since RPA-1 binds ssDNA, RPA-1 focus formation is dependent on DNA resection. Therefore, it is expected that MRE-11, the nuclease forming ssDNA, will be recruited to DSBs prior to RPA-1, suggesting that a time window of 15 minutes will be most appropriate for imaging MRE-11. Second, 15 minutes of total imaging permits data acquisition every 10 seconds, which allows high resolution of timing of focus formation (67% of foci appear in the first 2 min, Fig S1A) and allows the detection of MRE-11 foci with a duration of less than 2 minutes (15% of the foci). The only disadvantage of 15-minute imaging is that foci formed at later time points will not be detected (24% of foci, Fig S1B). To improve upon our data analysis utilized in the protocol from Koury et al., 2018, we use an automated focus calling software (for details see Materials and Methods). We examined recruitment kinetics of MRE-11 by 3D imaging every 10 seconds following microirradiation for 15 minutes to obtain a live time-course of MRE-11 recruitment (Fig 1). 1-day-old adults were microirradiated in 4 germline zones: pre-meiotic tip (PMT), transition zone (TZ), mid-pachytene (MP), and late-pachytene (LP). Images were analyzed using the FIJI Plug-In TrackMate to obtain number of foci per nucleus and the time of focus recruitment (see Materials and Methods).

We found that MRE-11 forms significantly more foci in MP compared to all other germline zones (average 2 foci/nucleus vs. 1 focus/nucleus; Fig 1C), similar to what we observed for RAD-51 (Koury et al., 2018). Another measurement for recruitment of MRE-11 to DSBs is the time it takes for a focus to appear. Although a similar number of sites may be eventually available for MRE-11, the speed by which a focus forms can vary. This could be indicative of the ability of MRE-11 to access damage sites. MRE-11 is recruited to microirradiation-induced breaks ∼2 minutes on average post-microirradiation in TZ and MP (early meiosis), which is significantly faster than the ∼3 minutes it takes for focus formation in PMT and LP (Fig 1D). The acquisition of RPA-1 recruitment time in (Koury et al., 2018) was done using different parameters (2-minute intervals). To directly compare recruitment of MRE-11 to RPA-1, we also analyzed MRE-11 at MP using 2-minute interval movies and the same focus size calling as in (Koury et al., 2018). By this analysis, MRE-11 appeared ∼4 minutes earlier than RPA-1 (Fig S1C). The time of recruitment is consistent with our previous data and the role of MRE-11 in generating ssDNA for recruitment of RPA-1 and RAD-51, both of which show average recruitment times longer than that of MRE-11. Altogether, this indicates that the ability of MRE-11 to be recruited to DSBs is different throughout the germline, and this difference may reflect regulation of its activity.

### Microirradiation induces similar levels of localized damage throughout the germline

We assume that the number of MRE-11 foci reflects the ability of MRE-11 to access the location of damaged DNA. However, this acts under the assumption that the same DNA damage was formed in each germline region tested. Since we used exactly the same laser power throughout the worm and all germline regions are positioned roughly the same distance from the laser and the edge of the worm’s body, we did not expect any differences in level of damage inflicted. To test this, we employed the terminal deoxynucleotidyl transferase-mediated dUTP nick-end labelling (TUNEL) assay, which marks free 3’-OH and in doing so marks DNA damage. The protocol utilized does not mark SPO-11 induced breaks (see Discussion). TUNEL is the most direct measurement of DNA damage created by laser microirradiation, as the mechanism of action of microirradiation is the formation of DNA nicks, and when clustered, nicks form DSBs (Lan et al., 2004). 1-day-old adults were exposed to microirradiation as indicated above. Microirradiated worms were then recovered and dissected 15 minutes post-microirradiation (Fig 2A). Laser microirradiated germlines exhibited localized TUNEL signal that was never found in non-microirradiated germlines, indicating the formation of free 3’-OH ends, as expected by laser microirradiation (Fig 2B, second row). The number of affected nuclei is consistent with DNA damage specific to microirradiated nuclei (Fig S2D).

**Figure 2.**
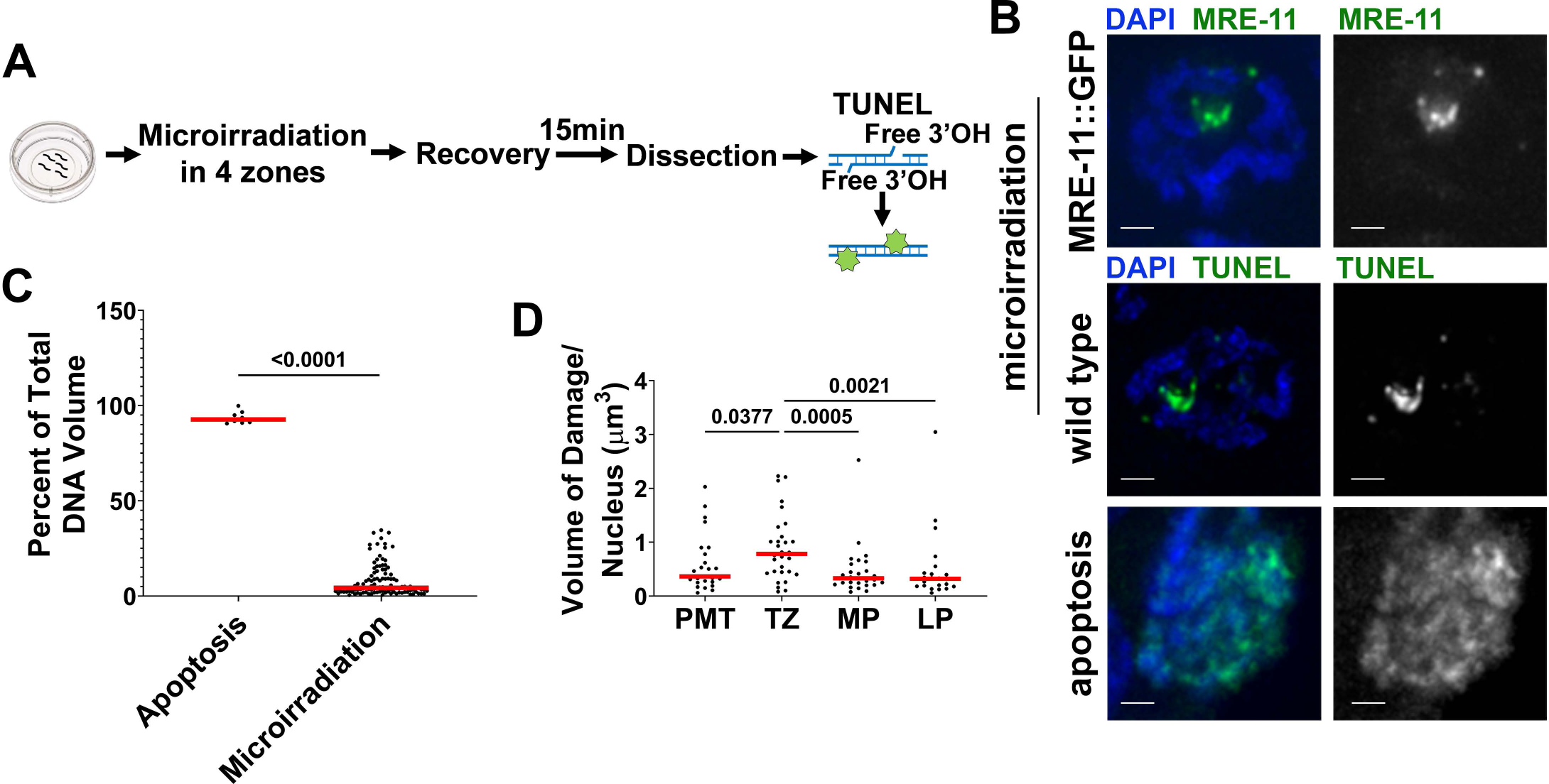
Microirradiation induces damage that is distinct from apoptotic levels of damage. **A)** Experimental design for the terminal deoxynucleotidyl transferase dUTP nick end labeling (TUNEL) assay. **B)** Examples of nuclei that were hit with microirradiation (top two rows) or are apoptotic (bottom row). The top row is an example of MRE-11::GFP seen after microirradiation in MP (no staining). The middle row is from a wild type worm and is an example of TUNEL staining in an MP microirradiated nucleus. The bottom row is TUNEL staining of an apoptotic nucleus in LP. TUNEL in the bottom row was performed in a *syp-1(me17)* mutant strain. Scale bars = 1µm. **C)** Percent of total DNA volume that is marked by TUNEL staining. The apoptosis column includes data from apoptotic nuclei in LP while the microirradiation column contains data from microirradiated nuclei in PMT, TZ, MP, and LP. Mann-Whitney U-test used to assess significance. **D)** Volume of damage per nucleus that is marked by TUNEL staining after microirradiation in each germline zone. Kruskal-Wallis was applied to determine significant differences between rank means, and if there was determination of significant differences Mann-Whitney U-test was applied for each pairwise comparison. Horizontal red lines on graphs in **C** and **D** indicate median.

Apoptosis also results in TUNEL positive nuclei due to fragmentation of chromosomes that is part of the apoptotic process. However, this type of DNA damage is not localized and affects all the nuclear DNA (Kressel & Groscurth, 1994). Apoptotic nuclei are present in high levels in LP nuclei of mutants that are impaired in SC formation (Alpi et al., 2003, MacQueen et al., 2002, Smolikov et al., 2009). To determine if microirradiation-induced DNA damage is distinct from that formed by apoptosis, the area of all stained regions through the image stack was taken and volume was calculated for each nucleus. The volume was then normalized to DAPI, to indicate the percent of DNA that contains 3’-OH DNA. This volume ratio was compared between microirradiated germline nuclei and apoptotic nuclei (Fig 2C, S2A). Microirradiated nuclei had on average only ∼2.9% of their DNA volume stained with TUNEL, compared to 93.6% in apoptotic nuclei (Fig 2C and D). This clearly indicates that the DNA damage following microirradiation is localized and distinctly different from apoptotic DNA damage.

Moreover, comparing the amount of damage between zones, as evidenced by the total area of staining per nucleus, indicated that the damage induced in each germline zone was similar, and only TZ nuclei exhibited an increase in damaged area compared to the other stages (Fig 2D). The organization of chromatin is changed as nuclei progress into and through meiosis, which is reflected in an increase in DAPI volume (Fig S2B). When normalized to DAPI volume, MP and LP showed a decrease in the relative area of TUNEL staining compared to other stages (Fig S2C). The percent of nuclei with TUNEL out of nuclei irradiated was consistent throughout the germline (Fig S2D). Overall, this determines that the damage induced in all of our studied germline regions was similar, localized, and not confounded by location within a whole worm. Most importantly, MP and LP were exposed to identical damage by all parameters tested (Fig 2 and S2), and showed identical protein expression (Fig 1B), indicating that the differences observed between these stages in their ability to recruit MRE-11 reflects a biological difference in MRE-11’s ability to access breaks.

### MRE-11 forms two types of focus configurations: individual foci and clusters

In our previous studies using fixed samples we noted that following microirradiation RAD-51 forms both individual foci and foci clusters (2 or more foci that are touching) (Koury et al., 2018). The size of foci in the “individual foci” category was identical to that of RAD-51 foci induced by SPO-11 that are considered sites in which a single DSB is formed (Fig S2F). This suggests microirradiation creates two classes of damage sites: ones that contain multiple events (clusters) and ones that likely contain individual events (individual foci). Analysis of fixed samples from MRE-11::GFP retrieved at different time points post-microirradiation (see below) was consistent with this observation (Fig 3A), indicating that both complex (clusters, 2 or more foci touching each other) and less complex (single focus) DNA damage is formed following microirradiation. These events were evenly split in proportion (Fig 3B), with minor differences between zones (also see below). The size of individual MRE-11 foci was indistinguishable from that of individual RAD-51 foci induced by SPO-11 or by microirradiation (Fig S2F). The resolution of the images obtained from live imaging analysis does not allow resolution of individual foci within a cluster (Fig 3A). However, such discrimination can be done by examining the size of the focus. We noticed that some foci formed were significantly larger than others. We used the average diameter of the clusters defined by fixed sample analysis to divide the categories of damage observed by live-imaging into clusters and foci (Fig 3A). Based on this analysis, we set the threshold to identify foci in the time course images as 0.6µm, and we used this threshold to distinguish between foci (<0.6µm) and clusters (>0.6µm). The ratio of these foci and clusters was then examined in each of the four zones microirradiated. This form of analysis yielded similar results to that obtained by fixed sample analysis: around 50% of damage regions formed upon microirradiation were clusters, and there was no significant difference between the four zones regarding ratio of clusters to foci (Fig 3C). Thus, our method of assigning foci versus cluster categories for live imaged foci likely reflects foci and clusters identified by immunofluorescence analysis. Altogether this data shows that microirradiation creates two types of DNA damage sites that vary in the amount of DNA damage. To refer to clusters and foci collectively we will use the terminology “recruitment regions”.

**Figure 3.**
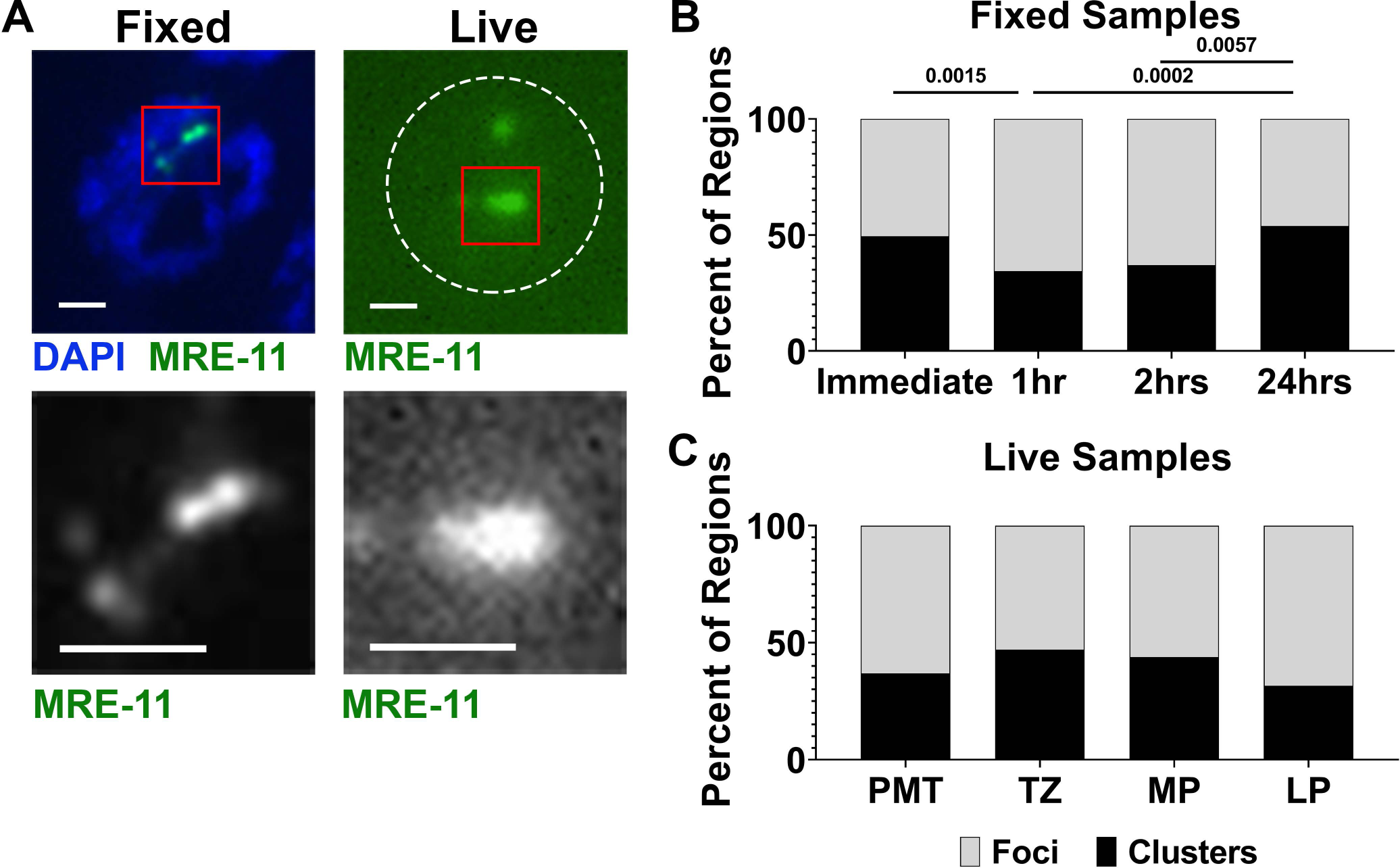
Clusters are made up of 2+ foci and can be resolved into individual foci in fixed images. **A)** Examples of clusters and foci in microirradiated nuclei that were fixed (left) or live imaged (right) MRE-11::GFP worms. Dotted white line is the nuclear outline in the right panel. Red square in each panel is the area that is in the close-up panel below. Scale bar = 1µm. **B)** Percent of total regions of DNA damage that are either foci or clusters in the fixed imaging. **C)** Percentage of microirradiation-induced recruitment regions that are clusters (>0.6µm, black) or foci (<0.6µm, gray) in the analyzed live imaging. Fisher’s Exact Test (two-tailed) was applied for all pairwise comparisons in **B** and **C**.

### MRE-11 shows an increase in recruitment over a 24-hour time period

Germline DSB repair occurs over the course of hours for both SPO-11-induced DSBs and microirradiation-induced DNA damage (Alpi et al., 2003, Koury et al., 2018). However, worms can endure live-imaging for no more than a couple of hours, which precludes the analysis of the whole process of DSB repair by live-imaging. To gain a better understanding of the duration and kinetics of repair of clustered DSBs in wild-type nuclei, we performed microirradiation, recovery, and gonad fixation at varying time points on our MRE-11::GFP strain. The drawback of this method is first, that the most immediate time point in which such analysis can be performed is 12 minutes following microirradiation, precluding analysis of early events of recruitment that live-imaging permits. Second, only nuclei with an MRE-11 focus or foci could be followed. The advantage of fixed sample analysis is two-fold. First, it provides the ability to look at the repair process hours and even days post-DNA damage induction. Second, it can provide more detailed information, as single foci can be resolved both separately and within clusters. As mentioned above, clusters were defined as 2 or more foci touching, with all other foci defined as separate foci (Fig 4A, green vs. red). Using this analysis, we could resolve the number of foci per cluster, and have an accurate estimate of the total number of foci in a nucleus (Fig 4A, black). We also classified a single cluster and a separate focus as a single “region” of DNA damage, to get an overall estimate of the amount of damage induced via microirradiation (Fig 4A, gray), providing a metric for comparison with our live-imaging data (the “recruitment regions”).

**Figure 4.**
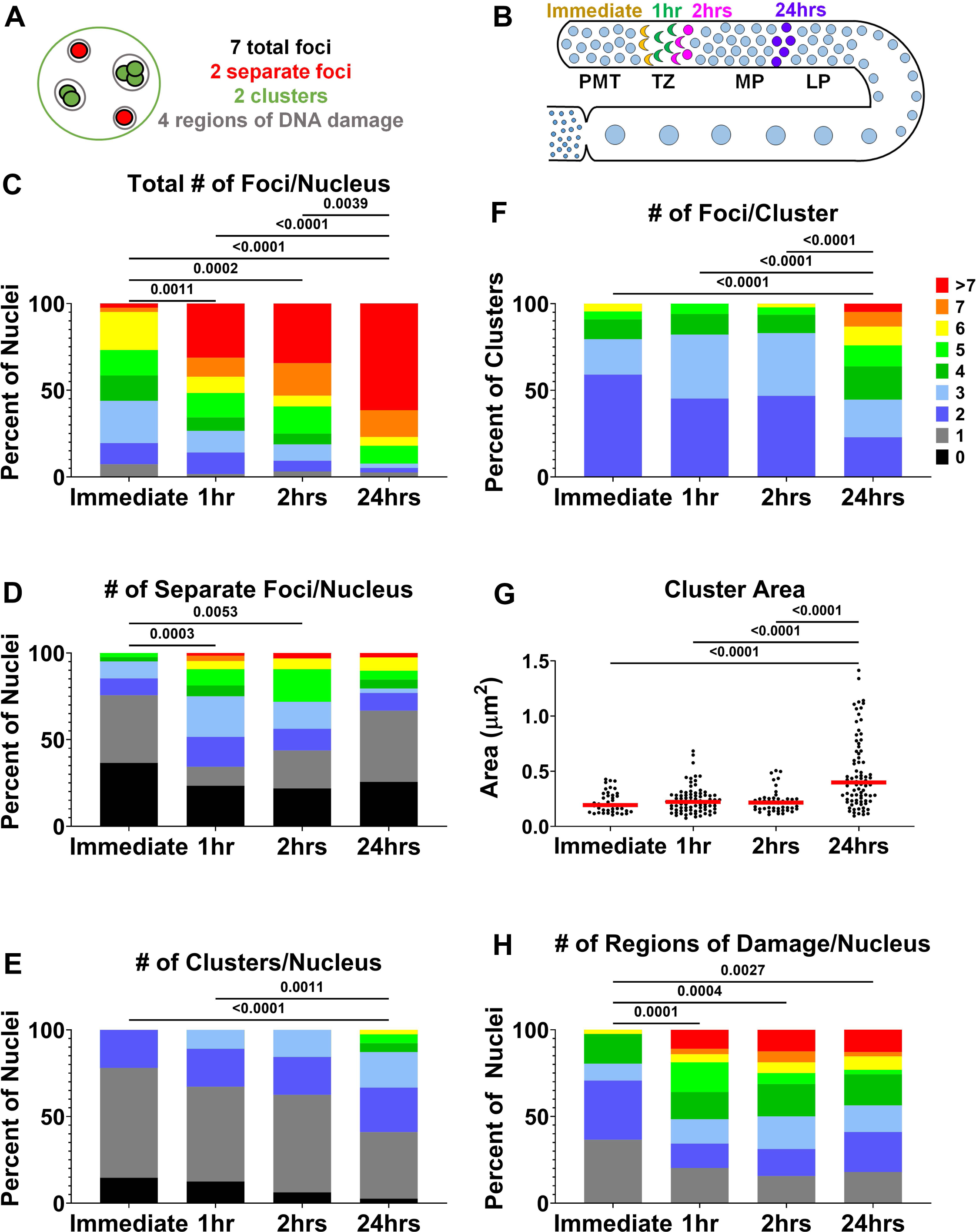
MRE-11 exhibits an increase in foci and clusters in a 24-hour time period following microirradiation. **A)** Cartoon depicting the terminology for classifying the microirradiation-induced damage in fixed samples. “Total foci” is the total number of foci in the nucleus (outlined in black). “Separate foci” are foci that are not touching any other foci (red). “Clusters” are 2 or more foci that are touching (green). “Regions of DNA damage” is the number of clusters and separate foci within the nucleus (outlined in gray). **B)** Schematic of the *C. elegans* germline. Worms are microirradiated in TZ and then EtOH fixed at indicated time points: Immediate (12 minutes), 1hr, 2hrs, or 24hrs. Each time point is marked with a distinct color in the germline cartoon and indicates where the nuclei are in the germline upon fixation at that time point. **C-F, H)** Legend to the right of figure **F** applies to all. **C)** Total number of foci (within clusters and separate foci) per nucleus. **D)** Number of separate foci per nucleus. **E)** Number of clusters per nucleus. **F)** Number of foci per cluster. **G)** Area of cluster (µm^2^). Each data point is a single cluster. Horizontal red line indicates median. **H)** Number of regions of DNA damage per nucleus. For data sets with more than 2 groups (**C-H**) Kruskal-Wallis was applied to determine significant differences between rank means, and if there was determination of significant differences Mann-Whitney U-test was applied for each pairwise comparison.

Microirradiation was performed in TZ, the region in which meiotic DSBs are typically formed. Four time points post-microirradiation were examined: immediate (12 minutes), 1, 2, and 24 hours post-microirradiation. Nuclei move in the germline at a rate of about 1 nucleus row per hour, thus the process of DSB repair occurs concurrently with progression through meiosis. The position of nuclei post-microirradiation was consistent with their rate of movement in the absence of microirradiation: at the immediate and 1 hour time points nuclei were still in TZ at the time of fixation, while nuclei examined at the 2 hour time point were in TZ transitioning to EP and nuclei at the 24 hour time point were either in MP or LP (Fig 4B).

MRE-11 shows a significant increase in the total number of foci per nucleus (this includes both separate foci and all foci present in clusters) over the time course analyzed (Fig 4C). There was an average of 4 foci per nucleus immediately after microirradiation, 6.2 1 hour post-microirradiation, 6.6 2 hours post-microirradiation, and 10.6 24 hours post-microirradiation. We then broke down the numbers of total foci per nucleus to assess whether separate foci or foci in clusters were contributing to this increase in the total number of foci. The number of separate foci per nucleus increased significantly between the immediate and 1-hour time points, and remained around 2-2.5 foci per nucleus for the remainder of the time course, potentially indicating that there is a maximum number of separate foci that will form (1 separate focus/nucleus at 1hr and 2 separate foci/nucleus at 2hrs; Fig 4D). The number of clusters per nucleus (2.1), the number of foci per cluster (4.1), and the area of each cluster (0.48 um^2^) is significantly higher at 24 hours post-microirradiation compared to all other time points analyzed (Fig 4E, 4F, and 4G). These data indicate that the significant increase in total foci between immediate and 1 hour time points is largely due to the appearance of separate foci, while the significant increase at 2 to 24 hours is largely due to expansion of foci number within clusters. The percentage of foci versus clusters reflects this trend as well, as there are significantly more foci than clusters in the 1 and 2-hour time points with respect to the 24 hour time point (Fig 3C). However, the total number of recruitment regions of DNA damage do not increase after the 1-hour time point (2.2 and 3.8 in immediate vs. 1 hour; Fig 4H). This could potentially mean that all sites of DNA damage have recruited MRE-11 to the point of detection by the 1 hour time point, and that following 1 hour, half of these sites continue to accumulate more MRE-11 over time.

Since nuclei move in the germline as time progresses, it is possible that the expansion of foci numbers from 2 to 24 hours is due to their movement from TZ to MP (when there is a lot of change in chromatin structure) and not due to the progression of repair. If so, microirradiation of nuclei in MP should lead to immediate cluster expansion (Fig S3). However, the number of foci per nuclei following microirradiation was similar in TZ and MP immediate time points and grows in similar proportion 1 hour post-microirradiation. Examining the structure of clusters at 24 hours post-microirradiation of MP nuclei was not attainable since these nuclei would have progressed to diakinesis. The overall number of foci per nucleus was higher in MP versus TZ, consistent with our live imaging data. Accumulation of MRE-11 foci within a cluster can be an outcome of recruitment of MRE-11 separate foci to an existing focus/cluster or expansion of a cluster. To test this, we performed hour long imaging of live worms (at lower intervals than the recruitment analysis to avoid bleaching). We observed no evidence for foci recruitment to clusters via live imaging (Fig S4). Altogether, this indicates that clusters increase foci numbers by measure of expansion, as a function of their progression in the process of DNA damage repair (see Discussion).

### Proteins from HR and NHEJ repair pathways show recruitment to, and colocalization at, microirradiation-induced DSBs

The formation of clusters indicates that about half of microirradiation regions contain several sites of DNA damage in very close proximity, indicative of complex DNA damage. Tissue culture cells exposed to high-linear energy transfer (LET) irradiation shift the repair pathway utilized from NHEJ to a combination of NHEJ and HR repair (Nickoloff et al., 2020). It could be interpreted in two ways: clustered DSBs are shifted towards HR, or clustered DSBs are repaired by mixed HR/NHEJ model. The germline is already committed to HR and NHEJ is only used when HR is inactive (Girard et al., 2018, Lemmens et al., 2013, Yin & Smolikove, 2013). In the condition of our experiment HR is fully active, thus involvement of NHEJ will suggest that clustered DSBs use mixed HR/NHEJ repair. To test this, we performed immunolocalization studies with co-staining for two repair proteins at a time at TZ 1 hour post-microirradiation using cKU-80 (a member of the KU complex) and MRE-11. We found that cKU-80 is recruited to microirradiation-induced breaks, and that it does show some colocalization with MRE-11 (MRE-11 with cKU-80 32%, and cKU-80 with MRE-11 24%, Fig 5A). The bulk of this colocalization occurs in clusters (Fig 5B and C). MRE-11 and cKU-80 colocalization in foci may reflect their joint function in some foci committed to an alternative-NHEJ (alt-NHEJ) pathway (see Discussion). Unlike MRE-11, RAD-51 activity is not found in any NHEJ pathway, thus cKU-80 and RAD-51 should only be colocalized in complex DNA damage. Indeed, cKU-80 and RAD-51 only colocalized within clusters and were not found colocalizing at foci (Fig 5D, E and F). We then examined colocalization between two HR proteins that act in different steps of repair: MRE-11 and RAD-51. Like we found above for other proteins, MRE-11 colocalized with RAD-51 in a fraction of cases (MRE-11 with RAD-51 ∼25%, and RAD-51 with MRE-11 ∼35%, Fig 5G), and clusters of both MRE-11 and RAD-51 exhibited more colocalization than did individual foci (Fig 5H and I). This is consistent with MRE-11 and RAD-51 representing two sequential steps in HR, and clusters containing multiple break events. To conclude, colocalization of DSB repair factors from opposing pathways/different steps in repair primarily occurs within clusters and not in separate foci. This suggests that clusters represent many breaks in one location that may be undergoing different forms of repair and that clustered DSBs recruit multiple repair pathways even in conditions that favor HR.

**Figure 5.**
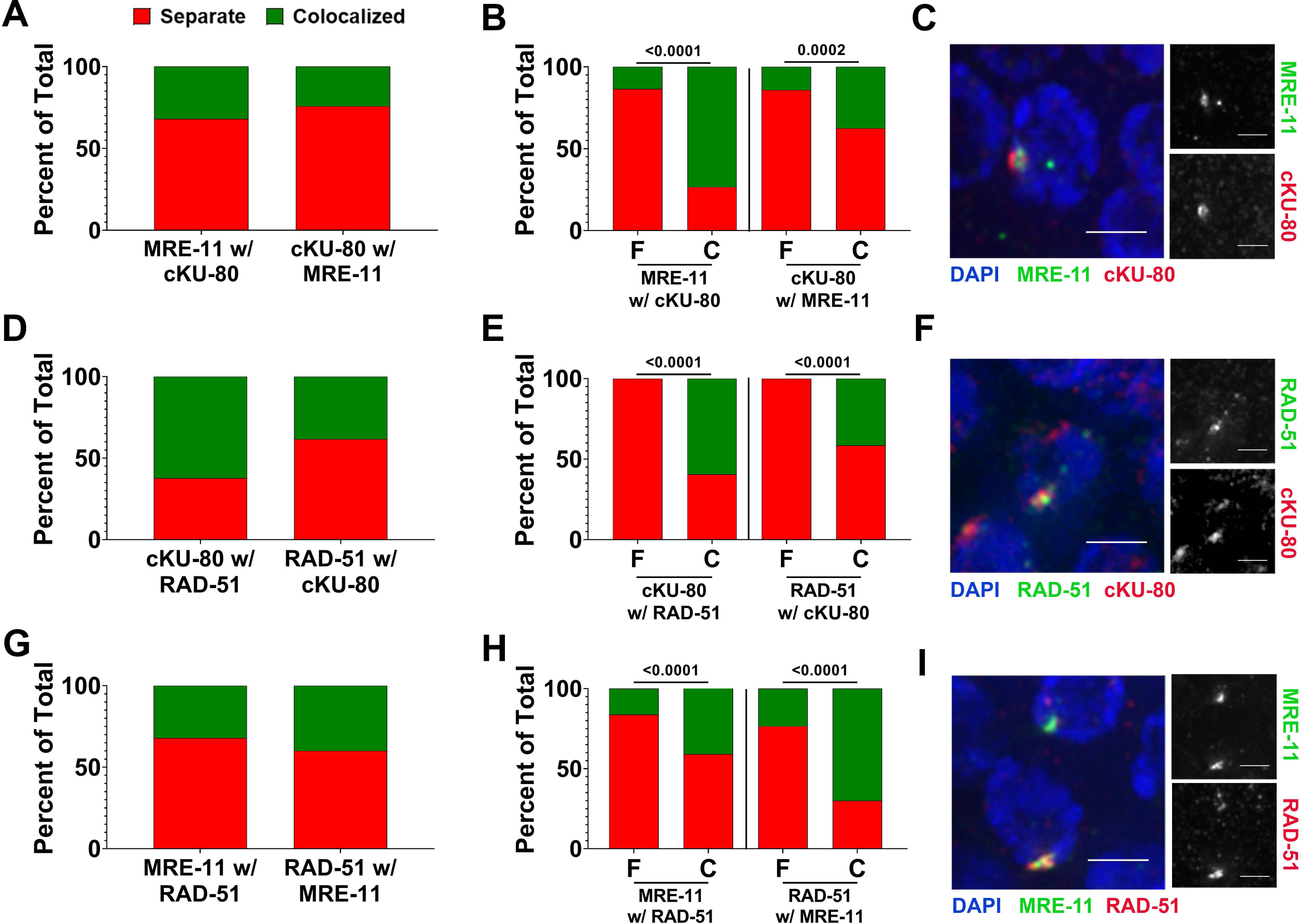
DNA repair factors colocalize in clusters more often than as foci 1hr following microirradiation in TZ. For classification of colocalization: separate indicates no overlap or touching (red), colocalized indicates overlap (green). **A-C)** Colocalization data for MRE-11 and cKU-80 marked with anti-OLLAS and anti-FLAG antibodies, respectively (*mre-11::ollas; flag::cku-80* strain). **A)** Colocalization for all types of microirradiation-induced damage (Foci and Clusters) for either MRE-11 with cKU-80 or for cKU-80 with MRE-11. **B)** Each data set in **A** split between foci (F) and clusters (C). **C)** Representative image of microirradiated nuclei after dissection and staining. Split channels on the right are MRE-11 (top) and cKU-80 (bottom). **D-F)** Colocalization data for cKU-80 and RAD-51 marked with anti-FLAG and anti-RAD-51 antibodies, respectively (*mre-11::ollas; flag::cku-80* strain). **D)** Colocalization for all types of microirradiation-induced damage (Foci and Clusters) for either cKU-80 with RAD-51 or for RAD-51 with cKU-80. **E)** Each data set in **D** split between foci (F) and clusters (C). **F)** Representative image of microirradiated nuclei after dissection and staining. Split channels on the right are RAD-51 (top) and cKU-80 (bottom). **G-I)** Colocalization data for MRE-11 and RAD-51 marked with GFP and an antibody against RAD-51, respectively (*mre-11::gfp* strain). **G)** Colocalization for all types of microirradiation-induced damage (Foci and Clusters) for either MRE-11 with RAD-51 or for RAD-51 with MRE-11. **H)** Each data set in **G** split between foci (F) and clusters (C). **I)** Representative image of microirradiated nuclei after dissection and staining. Split channels on the right are MRE-11 (top) and RAD-51 (bottom). For statistics performed in **B, E**, and **H**, a Fisher’s Exact Test was performed between the Separate and the Colocalized category. Scale bars = 3µm.

### Deletion of cKU-70 inhibits the formation of MRE-11 clusters in mitotic and early meiotic regions

With evidence that both NHEJ and HR factors are recruited to the DSB clusters induced by microirradiation, we wanted to determine whether recruitment of MRE-11 would be affected by the presence or absence of a functional NHEJ pathway. The kinetics of MRE-11 recruitment were largely unaffected by the deletion of cKU-70 (Fig 6), with a slight increase in the amount of time it takes for MRE-11 to appear in TZ (1.8 minutes for wild type vs. 2.5 minutes for *cku-70*; Fig 6B, middle panel) and an increase in the number of MRE-11 recruitment regions formed in LP (1 for wild type vs. 2.3 for *cku-70*; Fig 6D, left panel). The most notable difference observed with deletion of cKU-70 was the change in the percent of clusters and foci in PMT and TZ. In a *cku-70* background, MRE-11 forms significantly fewer clusters in PMT and TZ (36.9% in wild type vs. 10.2% in *cku-70* in PMT and 47.1% in wild type vs. 15.1% in *cku-70* for TZ; Fig 6A and B, right panel), whereas the relative proportion of clusters and foci was unchanged in MP and LP (Fig 5C and D, right panel). Since the overall number of foci is not increased in PMT, TZ, and MP, these changes must be attributed to a defect in MRE-11 recruitment to clusters, rather than clusters breaking apart to individual foci. A lower level of MRE-11 on clusters may lead to their classification as foci. Delay in recruitment may also explain the increase in recruitment time in TZ.

**Figure 6.**
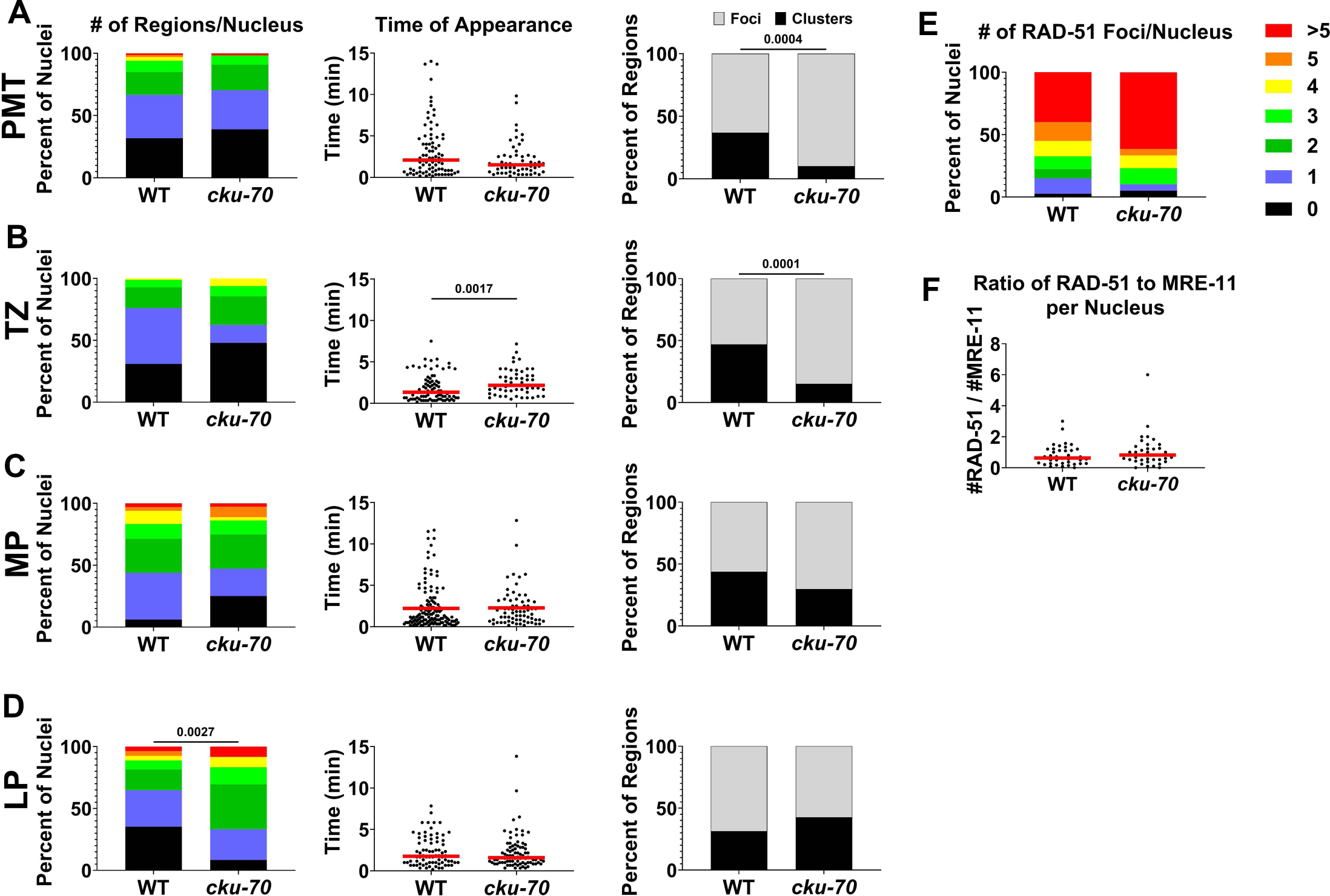
Deletion of cKU-70 leads to differential MRE-11 focus formation after microirradiation in PMT and TZ. *cku-70(tm1524)* mutant allele used for *cku-70* mutant in this figure. **A-D)** Far left column is the number of recruitment regions per nucleus in each germline zone, with each data point representing a single microirradiated nucleus. Figure legend in **E** applies to all graphs in the left column. Middle column is the time of focus appearance in minutes in each germline zone, with each data point representing a single recruitment region. Horizontal red bar indicates median. Far right column is the percentage of clusters (>0.6µm, black) versus foci (<0.6µm, gray) in each microirradiated zone. **E)** Number of RAD-51 foci per nucleus in nuclei microirradiated in TZ, recovered, and dissected 1hr post-microirradiation. **F)** The ratio of RAD-51 to MRE-11 foci within nuclei quantified in **E**, with the number of RAD-51 foci per nucleus divided by the number of MRE-11 foci within the same nucleus. Horizontal red bar indicates median. For **A-D** (left and middle columns) and **E-F**, Mann-Whitney U-test was employed. For the right column in **A-D** Fisher’s Exact Test (two-tailed) was used for all pairwise comparisons.

MRE-11 is required for resection of SPO-11 and gamma irradiation-induced DSBs in meiosis (Chin & Villeneuve, 2001, Yin & Smolikove, 2013). However, following microirradiation in TZ neither the number of RAD-51 foci per nucleus nor the ratio of RAD-51 foci to MRE-11 foci changed with abrogation of cKU-70 (Fig 6E and F). This is also reflected in the levels of colocalization of MRE-11 with RAD-51, which are similar between the two strains, and show a similar pattern of colocalization being largely between clusters as seen in Figure 5 (Fig S5A and B). This indicates that the impaired, but not abrogated, recruitment of MRE-11 is sufficient to promote resection (leading to RAD-51 focus formation) at clustered DSB damage sites.

### Deletion of COM-1 inhibits the recruitment and nucleolytic activity of MRE-11 in mid-to late-pachytene

MRN activity at SPO-11 breaks is dependent on COM-1/CtIP, its catalytic obligatory co-factor in meiosis (McKee & Kleckner, 1997). Indeed, *com-1* null mutants are phenotypically indistinguishable from *mre-11* catalytically null mutants (Penkner et al., 2007, Yin & Smolikove, 2013). Since MRE-11 is thought to be recruited to DSBs as part of a complex with SPO-11, COM-1, which does not play a role in DSB formation, is considered irrelevant to MRE-11 recruitment to DSB sites. However, microirradiation produces SPO-11-independent DSBs with no covalently bound proteins at the ends. This raises the possibility that COM-1 would play a different role in recruitment and activity of MRE-11 at these forms of damage. The previously described *com-1(t1626)* mutant was linked to a mutation that affected gonad length (*unc-32*), which complicates the comparable analysis to our other data performed in a wild type germline structure. Therefore, we generated a deletion mutant of COM-1 (*com-1(iow101)*) via CRISPR/Cas9 in an otherwise wild-type strain. *com-1(iow101)* contains an out-of-frame deletion which takes out 94% of the last exon (the most conserved region of COM-1), has a range of 1-12 DAPI bodies in diakinesis, and is homozygous sterile, all similar phenotypes to other COM-1 mutants that have been generated (Penkner et al., 2007).

COM-1 deletion did not have an effect on the timing of MRE-11 appearance in any of the germline regions (Fig 7A-D, right panel). However, there were significantly fewer MRE-11 recruitment regions that appeared in both MP (2.1 for wild type vs. 0.9 for *com-1*) and LP (1.4 for wild type vs. 0.5 for *com-1*; Fig 7C and D, left panel). Unlike what was found for *cku-70*, the ratio of clusters to foci was unchanged in all zones (Fig S5C-F). The significant change in the number of MRE-11 foci being recruited to microirradiation damage in MP and LP led us to test whether or not the activity of MRE-11 was similarly reduced in the later meiotic zones, as COM-1 in other organisms is shown to promote the nucleolytic activity of MRE-11. As a control, we also assayed the effect of COM-1 deletion on MRE-11 activity in TZ, a zone in which we saw no change in MRE-11 recruitment in our live imaging assay. Worms were microirradiated in TZ or MP, recovered, and then dissected and stained with an antibody against RAD-51 1 hour post-microirradiation. When microirradiated at TZ, no change in the levels of MRE-11 recruitment was observed. In terms of number of RAD-51 foci generated or in the ratio of RAD-51 to MRE-11 foci post-microirradiation, no difference was observed, suggesting that the deletion of COM-1 has no effect on the nucleolytic activity of MRE-11 at microirradiation-induced damage in TZ (Fig 7E and F). However, when the same experiment was performed in MP, significantly fewer RAD-51 foci were formed compared to the wild type background (11.6 RAD-51 foci per nucleus for wild type vs. 6.8 RAD-51 foci per nucleus for *com-1*; Fig 7G). Similar results were obtained at the immediate time point (12 minutes: 4.3 RAD-51 foci per nucleus for wild type vs. 1.6 RAD-51 foci per nucleus for *com-1*). The ratio of RAD-51 to MRE-11 was similarly reduced at both time points for MP in the *com-1* deletion mutants (Immediate: 0.4 for wild type vs. 0.2 for *com-1*; 1hr: 0.9 for wild type vs. 0.7 for *com-1*; Fig 7H). The colocalization of MRE-11 with RAD-51 was similar to wild type in both zones assayed (Fig 7I). Together, these data indicate that COM-1 enhances the recruitment and the nucleolytic activity of MRE-11 at complex DNA damage in specific stages of meiosis.

**Figure 7.**
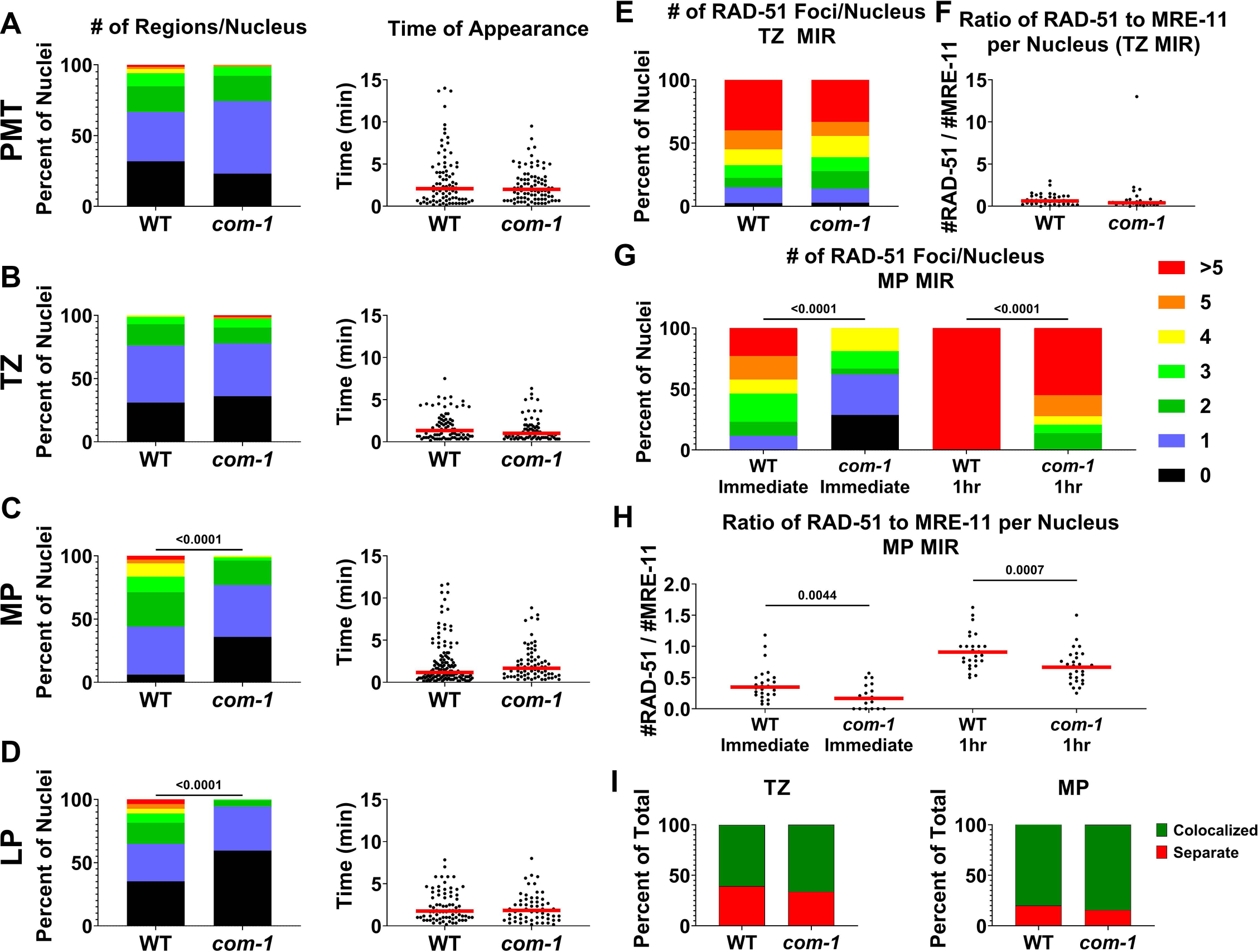
Deletion of COM-1 decreases MRE-11 recruitment and resection activity in MP and LP. *com-1(iow101)* mutant allele used in this figure for *com-1* mutant. **A-D)** Left column is the number of recruitment regions per nucleus in each germline zone. Right column is the time of focus appearance in minutes in each germline zone, with each data point representing a single focus. Figure legend in **G** applies to left column. **E)** Number of RAD-51 foci per nucleus in nuclei microirradiated in TZ, recovered, and dissected 1hr post-microirradiation. Figure legend in **G** applies here as well. **F)** The ratio of RAD-51 to MRE-11 foci within nuclei quantified in **E**, with the number of RAD-51 foci per nucleus divided by the number of MRE-11 foci within the same nucleus. **G and H)** Worms were microirradiated in MP, recovered, and dissected either immediately or 1hr later. **G)** Number of RAD-51 foci per nucleus. **H)** The ratio of RAD-51 to MRE-11 foci within nuclei quantified in **G**, with the number of RAD-51 foci per nucleus divided by the number of MRE-11 foci within the same nucleus. Each data point represents a single microirradiated nucleus. **I)** Colocalization of MRE-11 with RAD-51 in both wild type and *com-1* mutants microirradiated in either TZ (left) or MP (right) and dissected 1hr post-microirradiation. Horizontal red bar indicates the median in graphs in panels **A-D** (right column), **F**, and **H**. For **A-D** both columns, Mann-Whitney U-test was employed. For **E-H** Mann-Whitney U-test was used for statistical comparison between wild type and *com-1* mutants at both time points. For **I**, Fisher’s Exact Test was performed between Separate and Colocalized between wild type and *com-1* in each zone.

### The SC plays facilitating and inhibitory roles on recruitment of MRE-11 to microirradiation-induced DSBs

The SC is a meiosis specific complex that assembles between homologous chromosomes to target DSB repair to the homologous chromosome (as opposed to the sister chromatid) through the HR pathway. In the germline, the SC begins assembly in TZ, is fully formed in MP, and begins some disassembly at the end of LP (Nabeshima et al., 2005). The microirradiation performed at LP was in the nuclei just prior to SC disassembly. For both c*ku-70* and *com-1* mutants we saw effects that are different between pachytene regions (with SC) and PMT/TZ. We removed the SC using a *syp-3* mutant in the absence or presence of *cku-70. syp-3* encodes a structural protein of the SC (Smolikov et al., 2007b). Importantly, SYP-3 is a central region protein of the SC and has no role in DSB formation.

First, we tested if SC is playing a role in enabling or facilitating the formation of larger foci on clustered DSBs in the absence of *cku-70. syp-3* mutants had no effect on timing of MRE-11 recruitment or formation of clustered DSBs (Fig 8B and C). However, simultaneous deletion of both *syp-3* and *cku-70* led to a significant reduction in the relative proportion of clusters to foci in mid-pachytene (43.8% clusters (wild type) vs. 22.6% clusters (*cku-70; syp-3*); Fig 8C), indicating that both the SC and cKU-70 aid in MRE-11 recruitment in the germline to clustered DSBs. Defects in recruitment of MRE-11 may have an effect on its activity.

**Figure 8.**
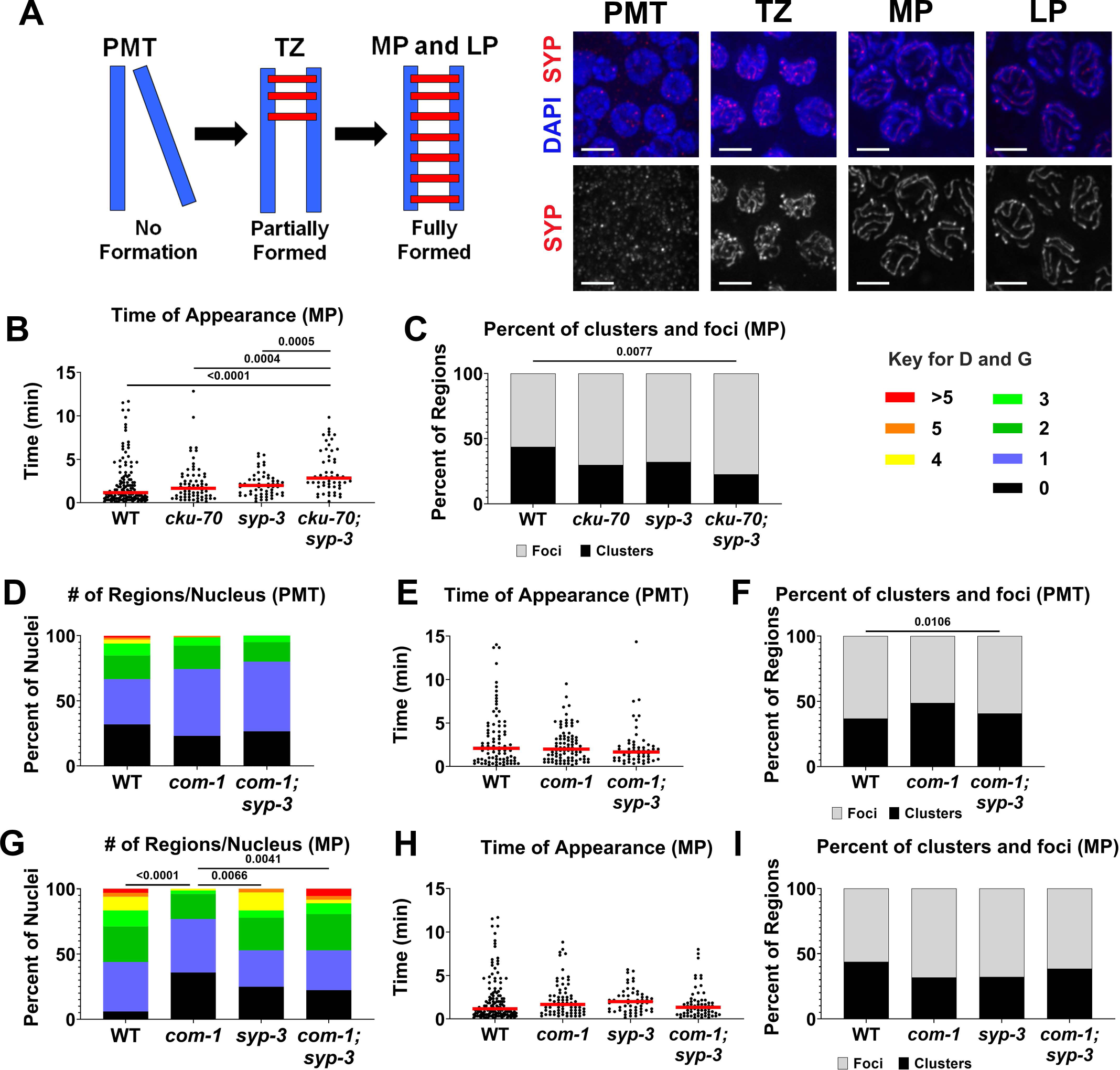
COM-1 is required for normal recruitment kinetics of MRE-11 in the presence of the synaptonemal complex. *syp-3(ok758)* mutant allele used in this figure. **A)** Progression of the formation of the synaptonemal complex (SC). In TZ synapsis is initiated, by MP the SC is fully formed, and at the end of LP there is partial disassembly of the SC. Blue bars in the cartoon represent homologous chromosomes and the red is the proteinaceous SC which forms between them. On the right are representative images of each germline zone with SYP-1 staining shown in red. Scale bar = 3µm. **B)** Time of appearance of regions of recruitment in minutes after microirradiation in MP. Each data point represents a single region. Horizontal red bar indicates median. **C)** Percentage of regions that are either clusters (>0.6um, black) or foci (<0.6um, gray) after microirradiation in MP. **D)** Number of recruitment regions per nucleus in PMT. **E)** Time of focus appearance (minutes) in microirradiated nuclei of PMT. Each data point represents an individual focus. Horizontal red bar indicates the median. **F)** The percentage of total recruitment regions in PMT that were either clusters (>0.6µm, black) or foci (<0.6µm, gray). **G)** Number of recruitment regions per nucleus in MP. **H)** Time of focus appearance (minutes) in microirradiated nuclei of MP. Each data point represents an individual focus. Horizontal red bar indicates the median. **I)** The percentage of total recruitment regions in MP that were either clusters (>0.6µm, black) or foci (<0.6µm, gray). Kruskal-Wallis was applied to determine significant differences between rank means in **B, D, E, G**, and **H**, and if there was determination of significant differences Mann-Whitney U-test was applied for each pairwise comparison. Fisher’s Exact Test (two-tailed) was applied for all pairwise comparisons in **C, F**, and **I**.

Data from the previous section indicates that COM-1 is aiding MRE-11 recruitment to, and function at, complex DSBs induced in the context of a fully formed SC. To test if the SC inhibits MRE-11 recruitment in the absence of COM-1, we tested MRE-11::GFP focus formation in *com-1*; *syp-3* double mutants. We performed microirradiation in PMT, a zone which has no SC formation in either background, and MP, a zone in which the SC is typically fully assembled. Simultaneous deletion of *com-1;syp-3* was no different in MRE-11 recruitment in PMT compared to either wild type or the *com-1* mutant (Fig 8D-F). However, when SC formation was abrogated in MP, *com-1;syp-3* mutants had significantly higher numbers of recruitment regions than the single *com-1* mutant and was similar to wild type (2.1, 0.9, 1.8 for wild type, *com-1*, and *com-1; syp-3* respectively; Fig 8G). Time of MRE-11 recruitment was unchanged in both mutants compared to wild type, as was the ratio of clusters to foci (Fig 8H and I). This suggests that COM-1 is required to stabilize the MRN complex in order to access damage induced in the context of a fully formed SC.

## Discussion

The germline is a tissue in which programmed DSBs are committed to repair by HR. Exogenously induced DSBs in meiosis can be targeted to HR, creating DSBs that can substitute for SPO-11 induced breaks, but may be also be repaired by other pathways. Here we have shown that clustered DSBs impose a different DSB repair pathway choice decision on meiotic nuclei that now recruit HR and NHEJ proteins. In our studies we analyzed the recruitment of MRE-11 to complex DNA damage and how it is modulated. We have shown that the recruitment of this protein to complex DSBs formed in meiosis is affected by the presence of the synaptonemal complex and NHEJ components. Our studies suggest that recruitment of DSB repair proteins to breaks can be a mode of regulation of their activity.

### Repair of clustered DSBs in the germline involves the recruitment of HR and NHEJ proteins

Ionizing radiation induces DNA damage in the form of nicks that when found adjacent to each other, form DSBs (Lan et al., 2004). Since nicks can be easily repaired, they are not considered mutagenic. However, DSBs are more deleterious and require repair prior to cell division. The mechanism of DSB formation is common to all forms of ionizing radiation including heavy particle radiation used in cancer therapy, gamma irradiation, and microirradiation (Nickoloff et al., 2020). However, these forms of radiation differ in the relative abundance of clustered DSBs, with high LET exposure (heavy ion) producing more clustered DSBs than low LET exposure (gamma irradiation). In agreement with studies from tissue culture, we show that microirradiation produces clusters of DSBs in the germline of intact organisms ((Koury et al., 2018) and this paper). This supports microirradiation as an effective method for the study of clustered DSB, and may explain why low LET exposure (gamma irradiation), cannot serve as a model for DNA damage equivalent to what is created by high LET radiation used in cancer therapy.

Studies in tissue culture cells indicated that the repair of clustered DSBs does not follow the normal repair program of the cell. Cells that typically engage in DSB repair via NHEJ, shift their repair pathway choice to a mixed HR/NHEJ model (Wang et al., 2010). In mammalian tissue culture studies NHEJ factors such as the KU complex accumulate at microirradiation-induced DNA damage, and whether or not the break undergoes NHEJ or HR, both pathways’ repair factors are often present (Aleksandrov et al., 2018, Mari et al., 2006, Muster et al., 2017). In the germline, DSBs are committed to repair via HR and we have shown that DSB clusters still recruit HR proteins ((Koury et al., 2018) and this paper). Since in tissue culture NHEJ shifted to an NHEJ/HR model, that may imply that HR is the preferred pathway for repair of clustered DSBs (Wang et al., 2010). In the germline, which is already committed to HR, this may suggest that repair of clustered DSBs may remain unchanged from the canonical HR program. However, we discovered that NHEJ proteins are recruited to microirradiation induced breaks, indicating that clustered DSBs recruit both NHEJ and HR proteins, regardless of the preferred repair pathway of the tissue exposed to irradiation. This indicates that clustered DSBs are a special form of DNA damage and unique repair environment that cannot be understood by directly extrapolating from studies of single DSB events.

The colocalization of MRE-11/RAD-51 and cKU-80 at clusters but not at foci indicates that HR and NHEJ events are targeted to different breaks. The partly overlapping presence of MRE-11, RAD-51, and cKU-80 at clusters suggests that some form of regulation is potentially occurring within these clusters to allow spatial organization of the repair proteins at designated sites. This could also mean that different forms of repair can occur in close proximity within one cluster. The requirement for multiple repair pathways in the repair of clustered DSBs may stem from the combination of different types of substrates at the break. Assuming a random positioning of nicks caused by microirradiation, it is likely that some will contain substrates favorable for c-NHEJ (blunt or almost blunt ends), while others can be targeted to MMEJ (short ssDNA overhangs) or HR (long ssDNA overhangs). Clustered DSBs also create new repair substrates, short pieces of dsDNA, that may be especially challenging to repair and thus attempt repair by several pathways (Pang et al., 2011). Studies from tissue culture point to defects in repair of such substrates by NHEJ, suggesting that recruitment does not necessarily mean repair on clustered DSBs.

### MRE-11 dynamics at complex DNA damage sites

MRE-11 is one of the first proteins recruited to exogenously-induced DNA damage in mammalian tissue culture cells, forming foci within the first few minutes following microirradiation-induced damage (Kim et al., 2005, Shamanna et al., 2016, Suhasini et al., 2013). Our studies identified similar kinetics in *C. elegans* germline nuclei, showing a remarkable evolutionary conservation of this trait in metazoans. The time by which most MRE-11 foci are appearing precedes that of RPA-1 and RAD-51 foci, in agreement with the role of MRN(X) in DSB resection.

As one of the first HR proteins recruited to the site of damage, and a protein required for efficient processing of DSB ends, the regulation of the activity of MRE-11 is expected to be a key point of DSB repair pathway choice. Here we have shown that MRE-11 kinetics at the sites of DSBs shows differences throughout meiotic prophase I. Alterations of accessory proteins (COM-1 or of competing pathways (cKU-70) changes these kinetics (also see below). MRE-11 forms more foci in MP than any other germline zone, similar to what we previously showed for RPA-1 and RAD-51 (Koury et al., 2018). Through the use of the TUNEL assay, we have shown that the level of DNA damage induced in the germline regions tested is similar, indicating that the differences are not due to increase in DNA damage in MP. Since MRE-11 action creates the ssDNA that RPA-1 and RAD-51 bind, the most parsimonious hypothesis will be that changes in MRE-11 recruitment lead to differential recruitment of RPA-1 and RAD-51. The enhanced recruitment of MRE-11 in MP may reflect upregulation of MRN activity in that meiotic stage or the presence of alternative and competing DSB processing mechanisms in the other germline regions. This differential regulation of recruitment shows that HR proteins can be regulated via their recruitment to microirradiation-induced DNA damage.

Studying MRE-11 recruitment kinetics also revealed that the structure of DNA damage sites changes over time. The greatest increase in regions of DNA damage happens between the immediate and 1-hour time points. This indicates that most DSB sites that can recruit MRE-11 have done so in the first hour (∼90% of sites in the first 30 minutes). On the other hand, the number of foci per cluster continues to increase through the time course. This cluster expansion could be an indication that repair is ongoing at these sites, and more MRN complexes are being recruited to this area over time. We found no evidence of fusion of foci in live imaging that we performed over a 1-hour time course, indicating that it is most likely expansion and not a fusion of separate foci together that is increasing the number of foci per cluster. The expansion of clusters may be attributed to chromatin relaxation. Alternatively, the accumulation of MRE-11 at these sites may reflect a conversion of local damage into substrates that are suitable for MRE-11 recruitment over time. Analysis of live imaging data indicates that MRE-11 is dynamic and further studies will be needed to determine if that reflects directional movement.

### KU at complex DNA damage

Following microirradiation, KU is found in both foci and clusters. Most cKU-80 foci do not contain HR proteins, indicating that it can associate with a unique class of DSBs. However, clusters that contain multiple DSBs can attract both HR and NHEJ factors on different DSBs within the same cluster (as discussed above). Although KU is recruited to MIR breaks it has a relatively small effect on MRE-11’s recruitment. We have shown that KU inhibits MRE-11 focus formation at microirradiation-induced damage at LP, but not at PMT, TZ, or MP. This may suggest that KU can compete with MRE-11 on binding to some DSBs formed in LP but not earlier. Interestingly, the formation of MRE-11 clusters is diminished in a KU deletion background in both PMT and TZ. If the reduction in clusters was due to breaking down of clusters, this should have been associated with an increase in overall regions of MRE-11 recruitment. Since the overall numbers of MRE-11 regions is not altered following KU removal, it is unlikely that the reduction of clusters reflects a breaking down of clusters. We favor the hypothesis that MRE-11’s ability to accumulate at clusters is reduced in KU mutants leading to the reclassification of the cluster as a focus. DSB clusters may be more susceptible to structural damage, as they contain short dsDNA fragments, and it is possible that KU’s action is required to maintain the integrity of other regions in the cluster, in a way that facilitates protein recruitment throughout the cluster. The synergistic effect of *syp-3* and *cku-70* mutants may support such a model, as SYP-3 is part of a complex with structural function at meiotic chromosomes. The effect of the SC on MRE-11 recruitment to complex DNA damage sites must be different than the one in DSB formation: the SC lateral element HTP-3 is involved in recruiting MRE-11 to chromosomes for DSB formation (Goodyer et al., 2008). But the lateral element still forms and DSBs are still created in SYP mutants (Smolikov et al., 2007a, Smolikov et al., 2007b). Taken together these results indicate that the SC and KU play redundant roles in MRE-11 localization to clusters, probably by maintaining the structural integrity of the chromosomes at the site of clustered DSBs.

### COM-1 at complex DNA damage

COM-1/Sae2/CtIP’s most conserved function is acting as an MRN(X) co-factor, promoting the resection activity of MRE-11 at the site of DSBs (Cannavo & Cejka, 2014, Clerici et al., 2005, Penkner et al., 2007). This activity is required for the removal of SPO-11 at meiotic DSBs by the MRN complex, especially in the presence of a functional NHEJ pathway (Clerici et al., 2005, Girard et al., 2018). COM-1 homologues have other more controversial roles: having their own endonuclease activity and/or DNA bridging functions. COM-1’s activity as an MRN co-factor is not completely clear and here we show that COM-1 is promoting the recruitment of MRE-11 to DSBs and, as a consequence, its resection activity. Since microirradiation induced breaks are likely available for all nucleases (not blocked by SPO-11), RAD-51 focus formation was not eliminated, only diminished when COM-1 was removed. The reason why MRE-11 recruitment to DSBs is inhibited only in MP and LP likely stems from the presence of the SC at these stages. In agreement, SC removal suppresses the defects found in COM-1 mutants in MP but has no effect on TZ nuclei. We propose that the presence of SC is an inhibiting factor for MRE-11 recruitment and is overcome by COM-1’s action in stabilization of the MRN complex. In agreement, a yeast point mutant in the SC axial component RED1, which abrogates SC formation, partially rescued the meiotic defect seen in *com1/sae2* mutants (Woltering et al., 2000). Taking together these findings with the ones above (interaction with KU), a complex picture emerges by which the SC is facilitating MRN-COM-1 complex recruitment to the chromosomes, but also has an inhibitory activity in the absence of COM-1.

Altogether these results support a model of regulation of MRE-11 at the level of recruitment to exogenously induced DNA damage. MRE-11 exhibits differing kinetics in a wild type background throughout meiotic prophase I, and this differential activity is modulated by alterations in other DNA repair pathways and accessory proteins. The complex damage induced by microirradiation also leads to the recruitment of other repair pathway proteins, namely KU, which forms both foci and clusters following damage induction. This presents a complex nature of regulation of pathway choice at exogenous DSBs in a meiotic context.

## Materials and Methods

### Strains Used

Worms were grown and maintained on nematode growth media plates that were seeded with *Escherichia coli* OP50. Plates were kept at 20°. All strains used were in the wild type N2 genetic background. The following strains were used:

N2 (wild type)

*mre-11(iow45[mre-11::gfp::3xflag])* V

*cku-70(tm1524)* III ; *mre-11 (iow45[mre-11::gfp::3xflag])* V

*cku-80(iow75[FLAG::cku-80])* III ; *mre-11 (iow95[mre-11::OLLAS])* V

*com-1(iow101)/ht2 [qls48]* I;III ; *mre-11 (iow45[mre-11::gfp::3xflag])* V

*com-1(iow101), syp-3(ok758)/ht2 [qls48]* I;III ; *mre-11 (iow45[mre-11::gfp::3xflag])* V

*syp-1(me17) V/nT1[unc-?(n754) let-? qIs50] (IV;V)*

### Genome editing via CRISPR/Cas9

The following strains were created via CRISPR/Cas9 injections: *mre-11 (iow95[mre-11::OLLAS])* V; *com-1(iow101)/ht2 [qls48]* I;III; *cku-80(iow75[FLAG::cku-80])* III. Worms were injected as 1-day old adults and recovered onto an NGM plate (for ssODN and crRNA sequences see Table S1). The following day, injected worms were singled. Singled plates were then screened for offspring showing rol or dpy phenotypes associated with mutation of dpy-10, the co-injection marker used in each CRISPR injection. Dpy, rol, and wild type siblings of dpy/rol phenotypes were then singled and screened for either an insertion or deletion, depending on injection performed. To verify that the tags did not disrupt the function of the protein we performed functional assays. KU loss of function has no effect on viability under normal conditions, but late embryos subjected to gamma irradiation show irradiation sensitivity (Clejan et al., 2006). Eggs were collected from FLAG::cKU-80 worms and subjected to gamma irradiation to determine sensitivity. Assessment of embryonic lethality and occurrence of abnormal progeny phenotypes revealed no significant difference from wild type (Table S2). Loss of function of *mre-11* leads to almost complete embryonic lethality due to inability to form or repair DSBs during meiosis (Chin & Villeneuve, 2001). MRE-11::GFP produced viable progeny indistinguishably from wild type (Table S3). These experiments altogether support the functionality of our tags.

### Microirradiation and Live Imaging

We followed the protocol outlined in (Harrell et al., 2018) for microirradiation of whole, live worms with the following modifications: for the experiments in all figures except Fig S1 and Fig S4 one worm was imaged at a time to allow for 10 second interval acquisition for 15 minutes without photobleaching. Z stacks of 10 images at 1µm intervals were taken at each time point. For data presented in Fig S1 images were taken every 2 minutes for 1 and a half hours and 2-3 worms were imaged at a time. For data presented in Fig S4 movies were taken at 30 second intervals for 1 hour, and two worms were imaged at a time. For data presented in Fig 2, 3B, 4, 5, 6E-F, 7E-I, S2, S3, and S5A-B and S5G-H: 5-10 worms were placed on a live imaging slide, microirradiated, and then recovered. For recovery following microirradiation 340µl of M9 was added onto the 10% agarose pad, allowing worms to be transferred to an NGM plate until dissection or fixation at desired time points.

### TrackMate Analysis

TrackMate v3.8.0 was used to track foci movement in the live images acquired in MetaMorph version 7.8.12.0 (Tinevez et al., 2017). TIF files were imported into FIJI as 8-bit files with a set scale of 1000 pixels in distance, known distance of 64.5 pixels, pixel aspect ratio of 1, and a unit length of micrometers. The frame was cropped to contain the microirradiated nuclei throughout all frames, then hyperstacked. Within TrackMate, the LoG Detector was used with an estimated focus size of 0.6µm (“blob size” in the program), which corresponds to the threshold of what we consider a small focus versus a cluster (see Results). All movies were blinded and threshold and quality filters for every movie were set to eliminate background. Hyperstack Displayer was used as the view. The Simple LAP Tracker was used with the following settings: frame to frame linking max distance of 1µm, max distance of 2µm, and a maximum frame gap of 5 frames. Filters on tracks were set for each movie to eliminate any background outside of the microirradiated nuclei. Linkage was performed by TrackMate with our parameters listed above, and subsequently double-checked with manual corrections when necessary. While connections between foci were manually checked, no additional foci were manually added within these tracks to avoid any biases.

### Intensity Measurements

One day old adult worms were dissected on charged slides and kept in the dark as much as possible to avoid bleaching. The following fixation protocol was used: soaking in -20^°^C methanol for 1 minute, 4% PFA for 20 minutes, a 10 minute 1xPBST wash, a 10 minute 4’,6-diamidino-2-phenylindole (DAPI, 1:10,000 of 5mg/ml stock in 1xPBST) wash, and a 1xPBST wash for 10 minutes-1 hour. Slides were imaged with MetaMorph version 7.8.12.0 with 100x/1.4 NA oil Leica illuminated with 110LED to capture whole gonad images. Intensity of light source was 5%, with a 500ms exposure time for both DAPI and GFP channels. 31 images were taken in a stack, with 0.2µm steps. Measurements of the intensity at the center plane of each nucleus were taken in FIJI in each zone corresponding to where microirradiation is normally performed. Fluorescent intensity was corrected to cytoplasmic background.

### MRE-11::GFP Time Course Analysis

One day old adult worms were microirradiated on live imaging slides in indicated zone (TZ (Figure 4) or MP (Figure S3)) and then recovered. Ethanol fixation was performed with the following protocol at indicated time points following microirradiation for each worm (as indicated in Results, Figure 4 and Figure S3): 5-10 adult worms were placed on an uncharged slide (Surgipath Leica) in 5µl M9 solution, most liquid was soaked up using filter paper, then 5µl ethanol was applied and allowed to evaporate, followed by application of 5µl M9-DAPI solution. Most of this solution was soaked up using filter paper again, and a coverslip with Vectashield was placed on top. Coverslips were sealed with nail polish. Images were taken on the DeltaVision wide-field fluorescence microscope (GE Lifesciences) with 100x/1.4 NA oil Olympus objective. Images were then deconvolved with softWoRx software (Applied Precision). Microirradiation was indicated by the presence of bright green foci, as no foci appear at endogenous DSBs, and these nuclei were scored.

### TUNEL Assay

Five, 1-day old adult worms were placed on a live imaging slide at a time according to the protocol outlined in Harrell et al., 2018. All four germline zones were targeted in both gonads of each worm and microirradiated at 15% attenuation (as for all microirradiation carried out in this paper). Worms were recovered and both gonads were dissected 15 minutes post-microirradiation. Terminal deoxynucleotidyl transferase-mediated dUTP nick-end labelling (TUNEL) protocol was followed as described in (Pauklin et al., 2009) with the following modification: slides were incubated for 30 minutes at room temperature in 0.1M Tris-HCl (pH 7.5) containing 1% BSA and no bovine serum. Imaging was performed on the DeltaVision wide-field fluorescence microscope as listed previously in MRE-11::GFP Time Course Analysis.

Volume of DNA, determined via volume of DAPI staining, was calculated using volume of a sphere equation (4/3πr^3^), with the inner, hollow sphere of the nucleus being subtracted from the outer sphere of DNA to get an accurate measurement of the DAPI volume:

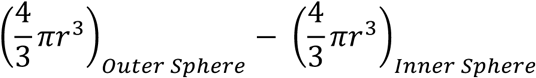

Radius of the sphere was calculated using known step sizes of 0.2µm between stacks and the top and bottom limits of the DAPI stain, and the area of the sphere was taken at the middle plane of the nucleus.

Volume of the TUNEL stain was determined via volume of a trapezoid, which employs the use of areas of irregular shapes. Each plane of the image was scrolled through for each nucleus and the total area of staining for every plane was calculated. The following formula was applied to determine overall volume:

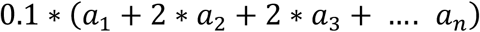

Where 0.2µm is the step size, therefore 0.1 is the radius of each plane. *a*_*1*_ is the area of the first plane of the nucleus with staining, followed by each subsequent plane (*a*_*2*_ and *a*_*3*_) until the final plane with staining is reached (*a*_*n*_).

### Immunostaining

One day old worms were microirradiated, recovered, and dissected in M9 solution on a coverslip and transferred onto positively charged slides, then placed onto metal blocks in dry ice. Worms were kept in the dark as much as possible throughout this protocol to avoid bleaching. Antibody staining was performed by the following procedure: wash in -20^°^C methanol for 1 minute, 4% PFA for 30 minutes, wash in 1xPBST for 10 minutes, block in 0.5% BSA in 1xPBST for 1-2 hours, then incubated with the primary antibody overnight at room temperature. Following primary antibody incubation, slides were washed in 1xPBST 1-3 times for 10 minutes each, then incubated with the secondary antibody for 2 hours at room temperature in the dark, followed by 1xPBST wash for 10 minutes, 10 minute staining in DAPI (1:10,000 of 5mg/ml stock in 1xPBST), and a wash in 1xPBST for 10 minutes-1 hour. All antibodies used were diluted in 1xPBST. For the SYP images presented in Figure 8A, no microirradiation was performed prior to dissection and staining. Primary antibodies used were: rabbit anti-RAD-51 (1:30,000), mouse anti-FLAG (1:500; Sigma F1804), rabbit anti-OLLAS (1:1,000; Genscript #A01658), goat anti-SYP-1 (1:500). Secondary antibodies used were: goat anti-rabbit Alexa Fluor 555 (1:500; Invitrogen), donkey anti-rabbit Alexa Fluor 488 (1:500; Thermo), and donkey anti-mouse Cy3 (1:500).

### Statistical Analysis

All statistics were performed using GraphPad Prism 8. For all data presented, a normality and logarithmic test was employed on the data to determine if the distribution of the data was normal or not. If all distributions were normal and there were only two groups being compared, a parametric t-test was employed. If all distributions were normal and there were more than two groups being compared, an ANOVA was employed to determine which groups had statistically significant differences and then a parametric t-test was performed to determine exact p-values for statistical differences. If even one distribution was not normal, and there were only two groups to be compared, Mann-Whitney U-test was performed. If there were more than two groups, Kruskal-Wallis was employed to determine significant differences between rank means. For all pairwise comparisons that presented with a significant difference through this test, Mann-Whitney U-test was performed to determine an exact p-value.

## Supporting information

Harrell et all 2020 Sup

## Acknowledgements

Some strains were provided by the *Caenorhabditis Genetics Center*, which is funded by the National Institutes of Health Office of Research Infrastructure Programs (P40 OD010440). We are grateful to A. Malkova, and members of the Smolikove laboratory for critical reading of this manuscript. We would like to thank D. Cooke, E. Koury, R. Lee and R. Bowman for their help with experimental procedures. This work was funded by National Institutes of Health (NIH) [R01GM112657 to S.S.]. Funding for open access charge: NIH [R01GM112657].

