## Supplementary material for "Recruitment of MRE-11 to complex DNA damage is modulated by meiosis-specific chromosome organization": Harrell et all 2020 Sup

Supplementary Figure 1

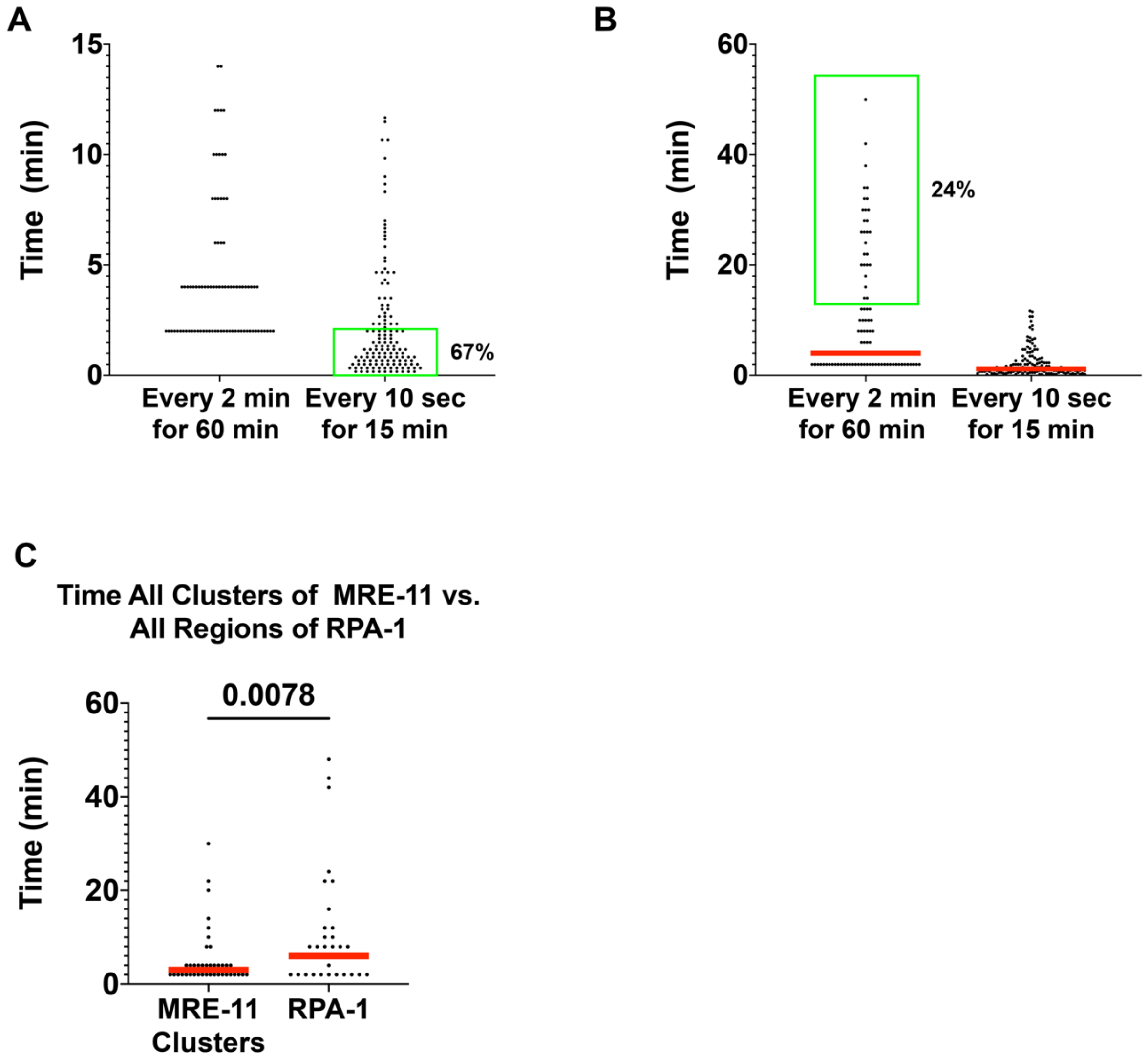


**Supplementary Figure 1. The majority of MRE-11 foci appear within 2min post-microirradiation, well before RPA-1. A)** Time of appearance (minutes) of MRE-11 from images taken every 2 minutes for one hour or every ten seconds for 15 minutes. Green box indicates the 67% of MRE-11 foci that appear in the first two minutes of imaging post-microirradiation. **B)** This is a close-up of the data presented in **A**, indicating the percentage of foci that appear after the first 15 minutes of imaging (24% of foci). **C)** Time of appearance in minutes of MRE-11 clusters and RPA-1 recruitment regions. MRE-11 clusters were used in order for an accurate comparison to RPA-1, given that the GFP::RPA-1 strain is so dim, the population of recruitment regions overrepresented is clusters. The horizontal red bar in panels **B** and **C** indicate median. Mann-Whitney U-test employed to determine significance.

**Supplementary Figure 2**

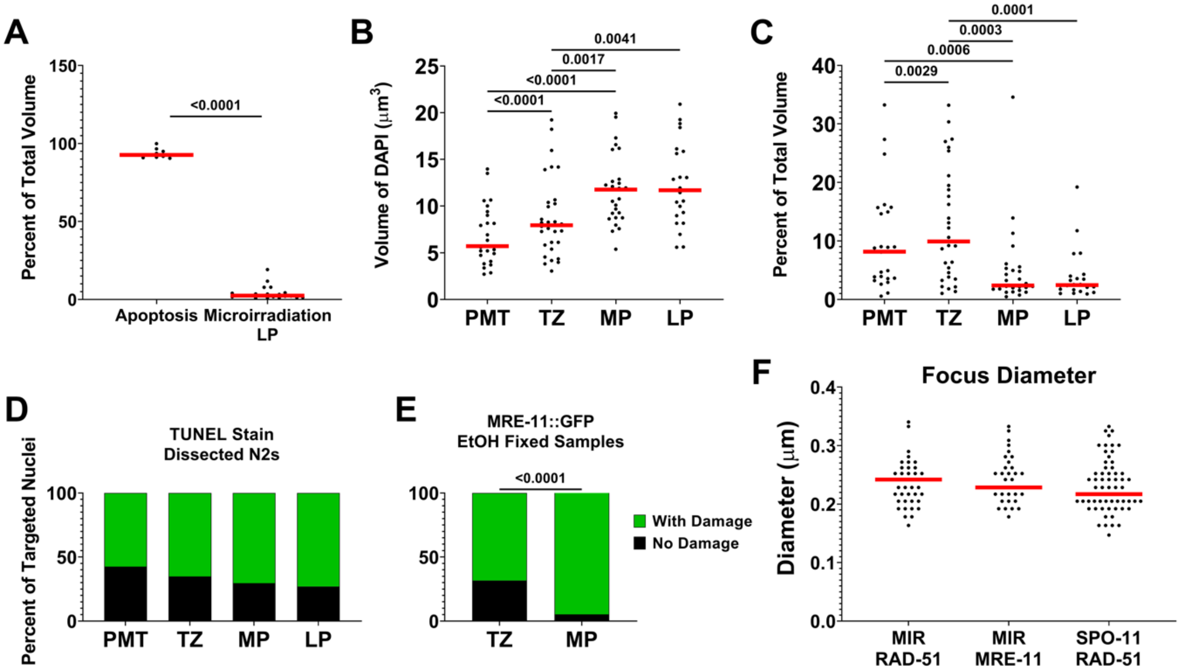


**Supplementary Figure 2. TUNEL staining and parameters tested for live imaging experiments. A)** Percent of total nuclear volume marked via TUNEL staining for nuclei undergoing apoptosis and nuclei microirradiated in late pachytene (LP). Each data point represents a single nucleus. Horizontal red bar indicates median. **B)** Total DNA volume per nucleus, marked via DAPI stain. Each data point represents a single nucleus. Horizontal red bar indicates median. **C)** Percent of total nuclear volume that is marked by TUNEL staining. Each data point represents a single microirradiated nucleus. Horizontal red bar indicates median. **D)** Percentage of targeted nuclei that were marked by TUNEL indicating presence of DNA damage (left) and percentage of targeted nuclei that had MRE-11::GFP foci in EtOH fixed samples (right). **F)** Diameter of individual foci induced via microirradiation (MIR) or SPO-11. Horizontal red bar indicates median. Kruskal-Wallis was applied to determine significant differences between rank means in **B, C, and F**, and if there was determination of significant differences Mann-Whitney U-test was applied for each pairwise comparison. Mann-Whitney U-test was also applied to data presented in **A** and **E.** Fisher’s Exact Test (two-tailed) was applied for all pairwise comparisons in **D.**

Supplementary Figure 3

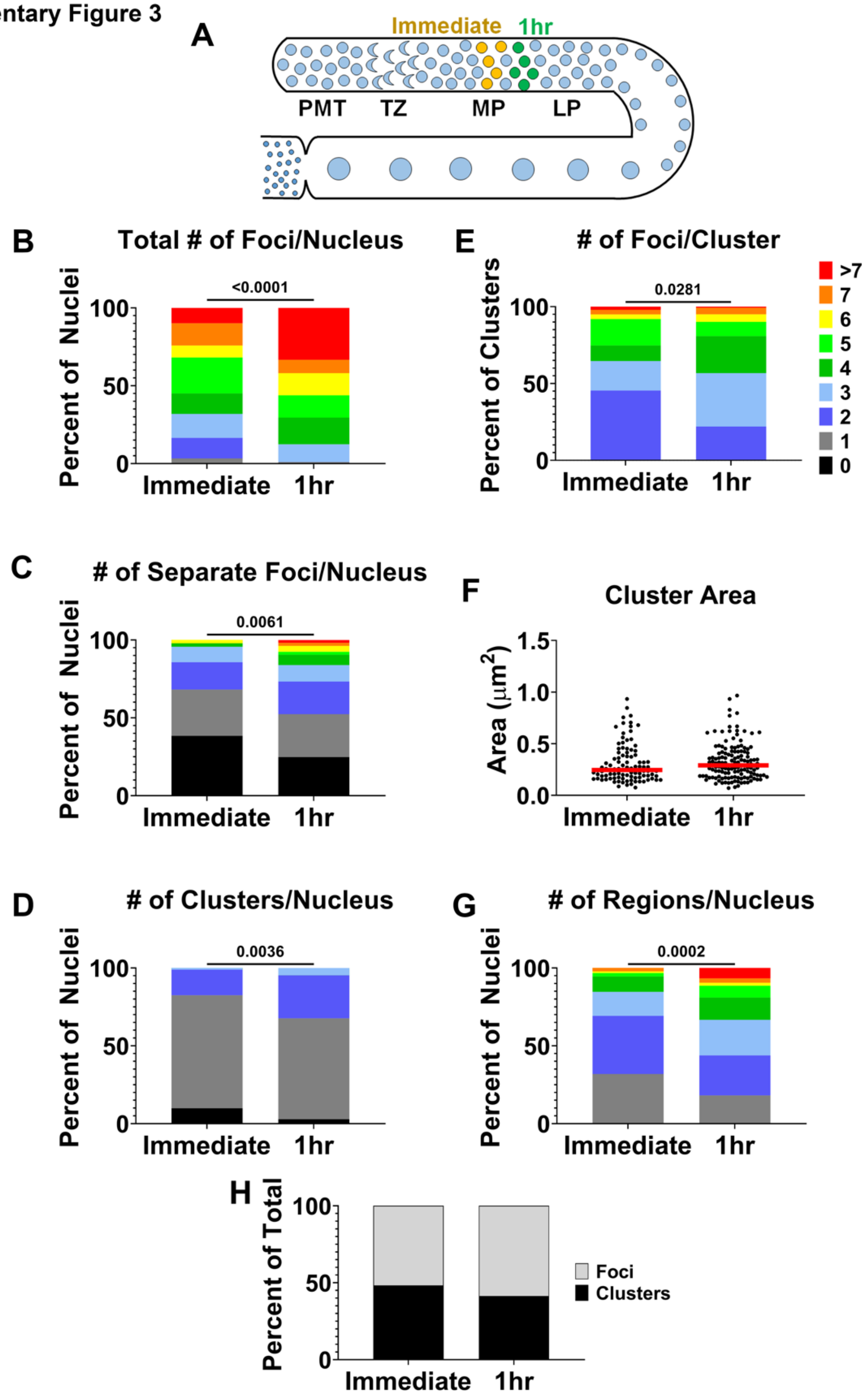


**Supplementary Figure 3. MRE-11 exhibits similar recruitment kinetics in nuclei microirradiated in MP as those measured in TZ following microirradiation. A)** Schematic of the *C. elegans* germline. Worms are microirradiated in MP and then EtOH fixed at indicated time points: Immediate (12 minutes) or 1hr post-microirradiation. Each time point is marked with a distinct color in the germline cartoon and indicates where the nuclei are in the germline upon fixation at that time point. **B-H)** See **Figure 4A** for classification of the microirradiation-induced DNA damage. The Figure legend in **E** applies to all graphs except for **F**. **B)** Total number of foci per nucleus. **C)** Number of separate foci per nucleus. **D)** Number of clusters per nucleus. **E)** Number of foci per cluster. **F)** Area of cluster ($\mu m^2$). Each data point represents a single cluster. Horizontal red bar indicates median. **G)** Number of regions of DNA damage per nucleus. **H)** The percent of total foci that are either in a cluster or separate (foci). Mann-Whitney U-test was applied for statistical analysis on **B-G**. Fisher's Exact Test used in **H.**

**Supplementary Figure 4**

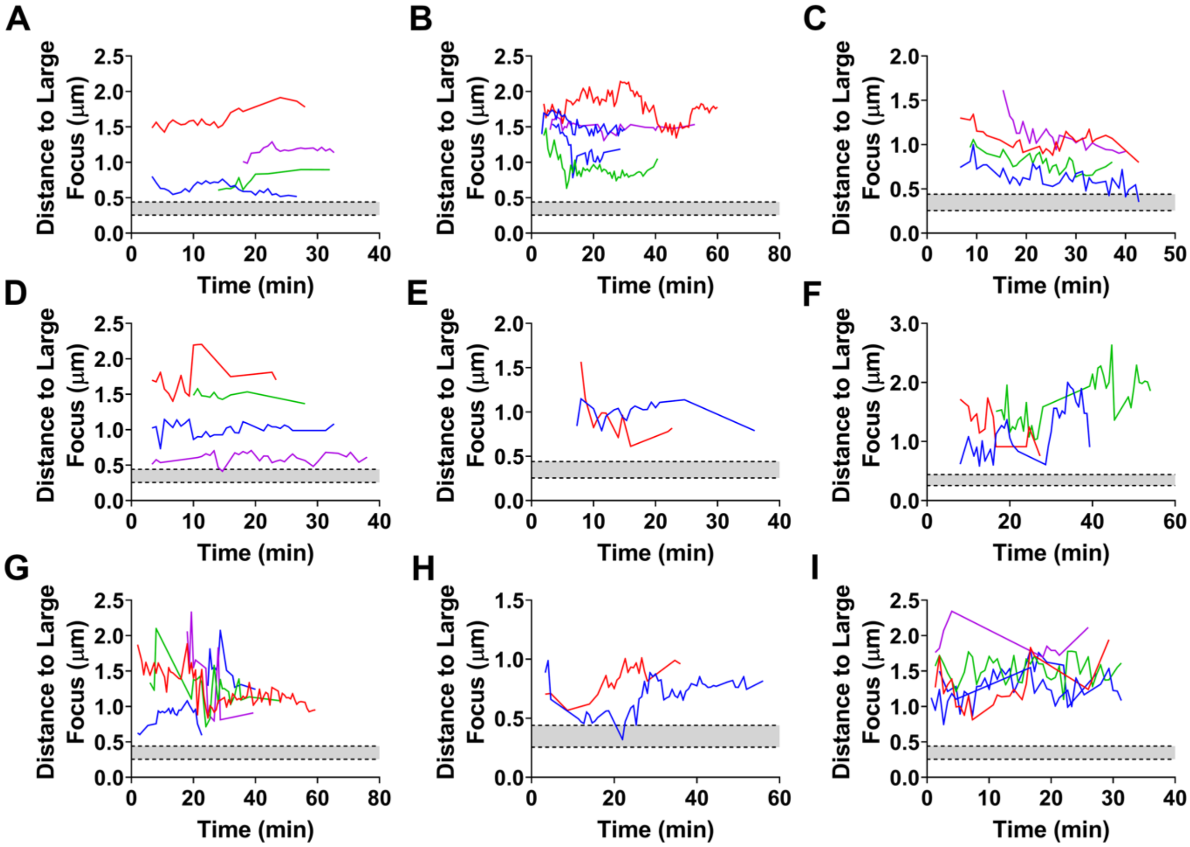


**Supplementary Figure 4. Foci do not show evidence of merging with clusters in a 1 hour time period. A-I)** Pairwise measurements between foci to clusters in each time frame that both are present over a 1 hour time period following microirradiation. The gray area indicates the average low (bottom dotted line) and high (top dotted line) diameter of each cluster in the nucleus analyzed (µm).

Supplementary Figure 5

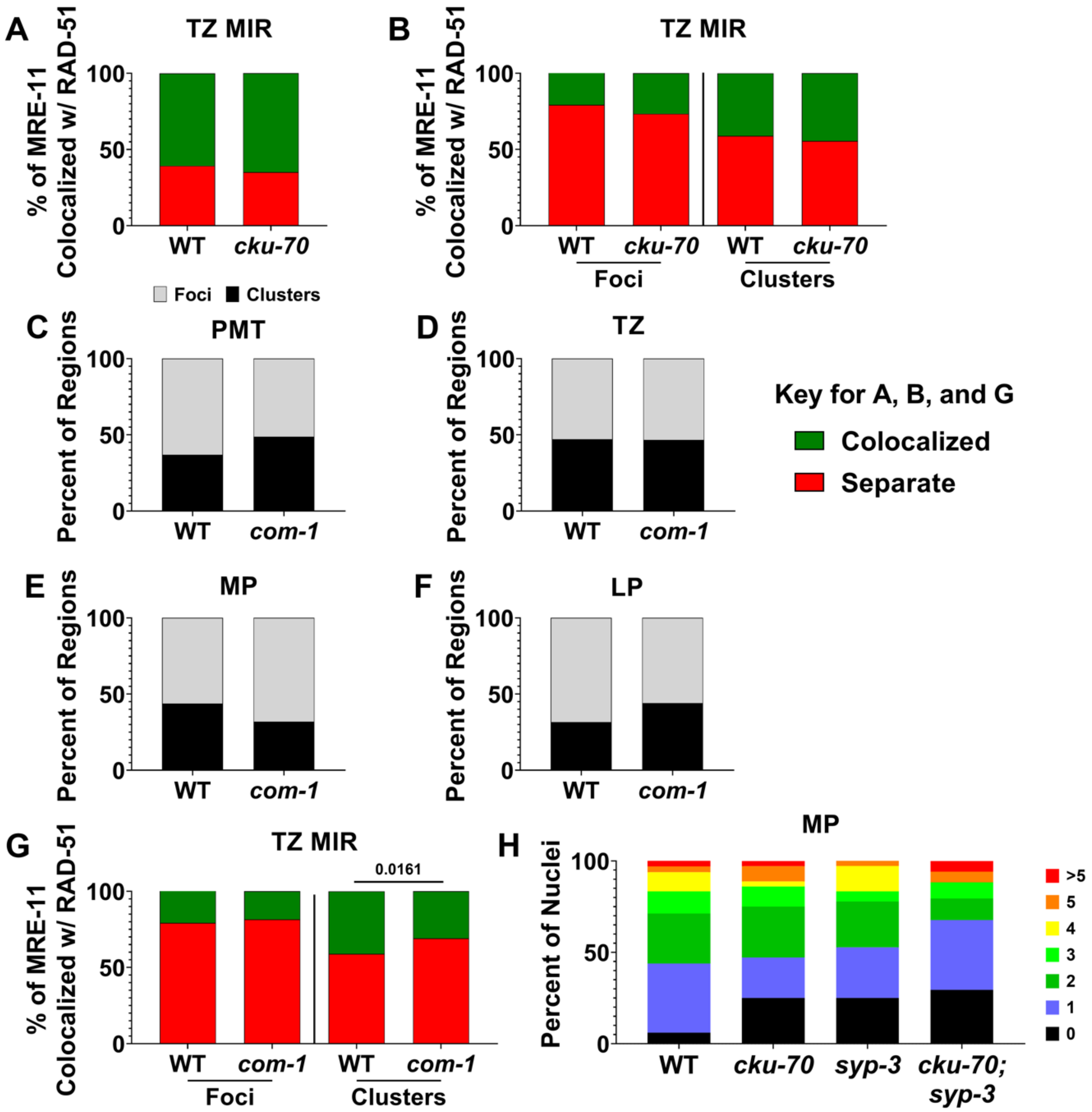


**Supplementary Figure 5. Breakdown of clusters and foci in live and fixed imaging following microirradiation. A)** Percent of colocalization of MRE-11 with RAD-51 following microirradiation in TZ for wild-type and *cku-70* mutants. **B)** Percent of MRE-11 colocalizing with RAD-51 split into foci and clusters for indicated genotypes. Same data as presented in **A**, only split here. **C-F)** Percentage of clusters (>0.6µm, black) versus foci (<0.6µm, gray) for wild type and *com-1* mutants in PMT **(C)**, TZ **(D)**, MP **(E)**, and LP **(F)**. **G)** Percent of colocalization of MRE-11 with RAD-51 following microirradiation in TZ for indicated genotypes split into foci and clusters. **H)** Number of recruitment regions per nucleus in worms microirradiated in MP. Kruskal-Wallis was applied in **H** to determine significant differences between rank means and no significant differences were found. Fisher's Exact Test (two-tailed) applied for all other data sets.

| **Sequence Name** | |
|---|---|
| *mre-11::OLLAS* ssODN | AGGAAAAGCAAGAGGAAAATCAGCTCCATCTAAGAAAAGGGATCTAAGTTTCTTCTCCGGATTCGCCAACGAGCTCGGACCACGTCTCATGGGAAAGTAAATAATTGTATTTTCACTTATCTCATTTACCGG |
| *mre-11::OLLAS* crRNA | GAAACUUAGAUCCCUUUUCU |
| *com-1(iow101)* crRNAs | GUAAAACUUCUCCCAUAAAU, GGCCGACUGGAAUCAAUACG |
| *FLAG::cku-80* ssODN | TCAACATCACCAATTTAAATTAAGTTACAGGAATGGACTACAAAGACCATGACGGTGATTATAAAGATCATGATATCGATTACAAGGATGACGATGACAAGCCGCCTAAAAAGGTATCTCCCGGGATTACCGTAAT |
| *FLAG::cku-80* crRNA | UACAGGAAUGCCGCCUAAAA |

**Table S1. crRNAs and ssODNs used for CRISPR/Cas9.**

| | **Avg. # Eggs Laid per P0** | **Avg. # F1 Adults per P0** | **Avg. %Progeny Viability** |
|---|---|---|---|
| **N2** | 266.3 ± 44.19 | 253.8 ± 35.71 | 95.6 ± 3.1% |
| ***mre-11::gfp*** | 296.7 ± 38.83 | 287.5 ± 39.85 | 96.8 ± 2.6% |

**Table S2. Fecundity Assay.** No significant differences detected with Mann-Whitney U-test. ±SD.

| | Avg. # Adults Hatched | Avg. % Adults w/ Abnormal Phenotype (Pvl, Vul, Unc) |
|---|---|---|
| **N2** | 66.2 ± 32.54 | 32.7 ± 15.32% |
| ***cku-70*** | 5 ± 8.04* | 91.2 ± 37.55% |
| ***FLAG::cku-80*** | 46.2 ± 15.17 | 39.2 ± 14.72% |

**Table S3. Gamma Irradiation Hypersensitivity Assay.** * = significantly different from N2 and *FLAG::cku-80* (less than 0.03) with Mann-Whitney U-test. No other significant differences in the table. ±SD.
